# Hormone-induced enhancer assembly requires an optimal level of hormone receptor multivalent interactions

**DOI:** 10.1101/2022.10.28.514297

**Authors:** Lizhen Chen, Zhao Zhang, Qinyu Han, Leticia Rodrigues, Emily Zboril, Rashmi Adhikari, Xin Li, Su-Hyuk Ko, Pengya Xue, Emilie Smith, Kexin Xu, Qianben Wang, Tim Hui-Ming Huang, Shasha Chong, Zhijie Liu

## Abstract

Transcription factors (TFs) activate enhancers to drive cell-specific gene programs in response to signals, but our understanding of enhancer assembly during signaling events is incomplete. Here, we show that Androgen Receptor (AR), a steroid hormone-regulated transcription factor, forms condensates through multivalent interactions in response to androgen signaling to orchestrate enhancer assembly. We demonstrate that the intrinsically disordered N-terminal domain (NTD) of AR drives 1,6-Hexanediol-sensitive condensate formation and that NTD deletion or aromatic residue mutation reduces AR self-association and abolishes AR transcriptional activity. AR NTD can be substituted by intrinsically disordered regions (IDRs) from selective proteins for AR condensation capacity and transactivation function. Surprisingly, strengthened AR condensation capacity caused by extending the polyQ tract within AR NTD also leads to impaired transcriptional activity without affecting AR binding on enhancers. Furthermore, either NTD deletion or polyQ extension reduces heterotypic multivalent interactions between AR and other enhancer components. These results suggest the importance of an optimal level of AR condensation in mediating AR-AR homotypic and AR-cofactor heterotypic interactions to regulate enhancer assembly in response to signals. Our study supports the notion that alteration of the fine-tuned multivalent IDR-IDR interactions might underlie AR-related human pathologies, thereby providing novel molecular insights for potential therapeutic strategies to treat prostate cancer and other AR-involved diseases by targeting AR multivalent interactions.

## Introduction

Androgen receptor (AR) is a member of the nuclear receptor superfamily that regulates transcription in response to steroid hormones. AR is involved in cell growth and survival and is known as the main driver of prostate cancer development, progression and therapy resistance. AR pathway has been the primary therapeutic target in advanced prostate cancer (Tan et al., 2015). Another AR-related human disorder is Kennedy’s disease, which is associated with unfolded protein response and transcriptional dysregulation caused by AR polyQ expansion (McCampbell et al., 2001; McCampbell et al., 2000; Thomas et al., 2005). In the absence of androgens, AR is bound by Heat Shock Protein 90 (HSP90) in the cytosol. When AR binds to androgens such as testosterone or dihydrotestosterone (DHT), AR undergoes conformational changes and is released from HSP90, resulting in the exposure of the nuclear localization signal (NLS) and nuclear translocation (Tan et al., 2015). Once in the nucleus, AR dimers bind to androgen response elements (AREs), which are predominantly located at distal enhancers (Stelloo et al., 2019).

Enhancers are *cis*-regulatory elements, which are found mostly in the intergenic and intronic regions, with well-established roles in regulating cell type-specific gene expression (Plank and Dean, 2014). Enhancers consist of clusters of transcription factor binding sites (TFBS) and are bound by a wide variety of transcription factors (TFs), coregulators, chromatin structuring and remodeling factors, RNA pol II, and other enzymes upon activation. Through a large-scale protein assembly, the enhancer complex loops over to interact with the target promoter to regulate transcription (Panigrahi and O’Malley, 2021). We have previously revealed dynamic high-order TF-DNA, TF-TF, and TF-cofactor interactions as a mechanism to regulate enhancer assembly under normal conditions and in the contexts of disease progression (Bi et al., 2020; Liu et al., 2014; Zhu et al., 2019). However, the precise mechanisms underlying signal-induced enhancer assembly remains elusive. Activation domains (ADs) of TFs are generally intrinsically disordered in the amino acid sequences and other cofactors including MED1 and BRD4 also mostly contain intrinsically disordered regions (IDRs) (Boija et al., 2018; Sabari et al., 2018). Recently, there has been an explosion of discoveries on how intrinsically disordered proteins (IDPs) undergo multivalent, dynamic but specific interactions and contribute to enhancer function by forming “hubs” or “condensates”, which can develop into phase separation systems under certain conditions (Boehning et al., 2018; Cho et al., 2018; Chong et al., 2018; Kwon et al., 2013; Lu et al., 2018; Sabari et al., 2018). While these hubs/condensates have been found to allow the IDPs to form strong and specific interactions to achieve enhancer regulation, new insights into how IDR-mediated multivalent interactions mediate signal-induced enhancer assembly would significantly advance our understanding of enhancer regulation.

Like other nuclear receptors, AR comprises an intrinsically disordered N-terminal domain (NTD) that is the transcriptional activation domain, a central DNA binding domain (DBD) with two zinc finger motifs, and a C-terminal ligand-binding domain (LBD). It has been previously reported that AR recruits a large number of coregulators, including coactivators (e.g., mediator complex), ATP-dependent chromatin remodeling proteins (e.g., SWI/SNF-BRG1), and pioneer factors (e.g., FOXA1) to enhancers in response to androgen signaling (Russo et al., 2019; Stelloo et al., 2018). However, it is not clear how AR assembles active enhancers upon hormone stimulation and whether IDR-mediated multivalent interactions play a role in hormone-induced enhancer assembly.

In this study, we demonstrate that the intrinsically disordered NTD of AR can undergo multivalent interactions and drive AR condensate formation. In androgen-sensitive cells, androgen induces AR condensation coupled with AR nuclear translocation. Through genetic and chemical approaches, AR condensation through its multivalent interactions can be weakened or strengthened. Our genomic and proteomic studies demonstrate that AR mutants with either weakened or strengthened condensation propensity have deficiencies in hormone-induced enhancer assembly and transcriptional activation. Our quantitative LacO array imaging data further confirmed that these AR mutants could not maintain normal homotypic AR-AR interaction and heterotypic AR-MED1 interaction. These results suggest that an optimal level of AR multivalent interactions is required for its proper function and that alterations of the fine-tuned AR condensation behavior might underlie human pathologies. Our results provide novel molecular insights for potential therapeutic strategies to treat prostate cancer and other AR-involved diseases by targeting AR multivalent interactions.

## Results

### AR NTD is an intrinsically disordered region and can drive phase separation in response to hormone stimulation

Phase-separated condensation mediated by intrinsically disordered regions (IDRs) of enhancer components has emerged as a novel mechanism of transcriptional regulation (Boija et al., 2018; Cho et al., 2018; Nair et al., 2019; Sabari et al., 2018). Since AR N-terminal domain (ARNTD) contains amino acid sequences predictive of an IDR (Figures S1A, 1A, and 1B), we hypothesize that ARNTD can drive phase separation. To test this hypothesis, we first used the optogenetic system, optoDroplet, which is based on *Arabidopsis thaliana* CRY2 (Bugaj et al., 2013). We expressed ARNTD (amino acids 1-538, interchangeable with IDR throughout the study) that was fused to mCherry and the photolyase domain of the CRY2 protein in 293T cells (Figure 1C). Upon blue light illumination, CRY2 undergoes self-association, leading to an increase of local concentration of the fused protein (Shin et al., 2017). Consistent with previous reports, CRY2-mCherry alone showed homogeneous distribution in the cell upon blue light activation (Figure 1D). Interestingly, fusing ARNTD to CRY2 lead to blue-light dependent formation of large, discrete, and droplet-like puncta. In contrast, C-terminus of AR (AR∆IDR) fused to CRY2 did not form droplet-like puncta (Figure 1D). Fluorescence recovery after photobleaching (FRAP) that reflects molecular dynamics (diffusion and binding) has been widely used to assess the liquidity of phase separation systems (Alberti et al., 2019). We performed FRAP experiments on the blue light-induced ARNTD optoDroplets and observed a slow and partial recovery (29 ± 9.3% recovery) (Figure 1E), suggesting that these puncta exhibit partially gel-like character (Shin et al., 2017). We next purified recombinant GFP-ARNTD protein (Figures S1B and S1C), and performed *in vitro* droplet formation assays. Purified GFP-ARNTD fusion protein formed spherical droplets in the presence of PEG8000 as crowding agent and the droplet size correlated with protein concentration (Figures 1F and 1G). Together, the live-cell optoDroplet and *in vitro* droplet formation assays suggest that AR NTD is able to undergo phase separation.

**Figure 1.**
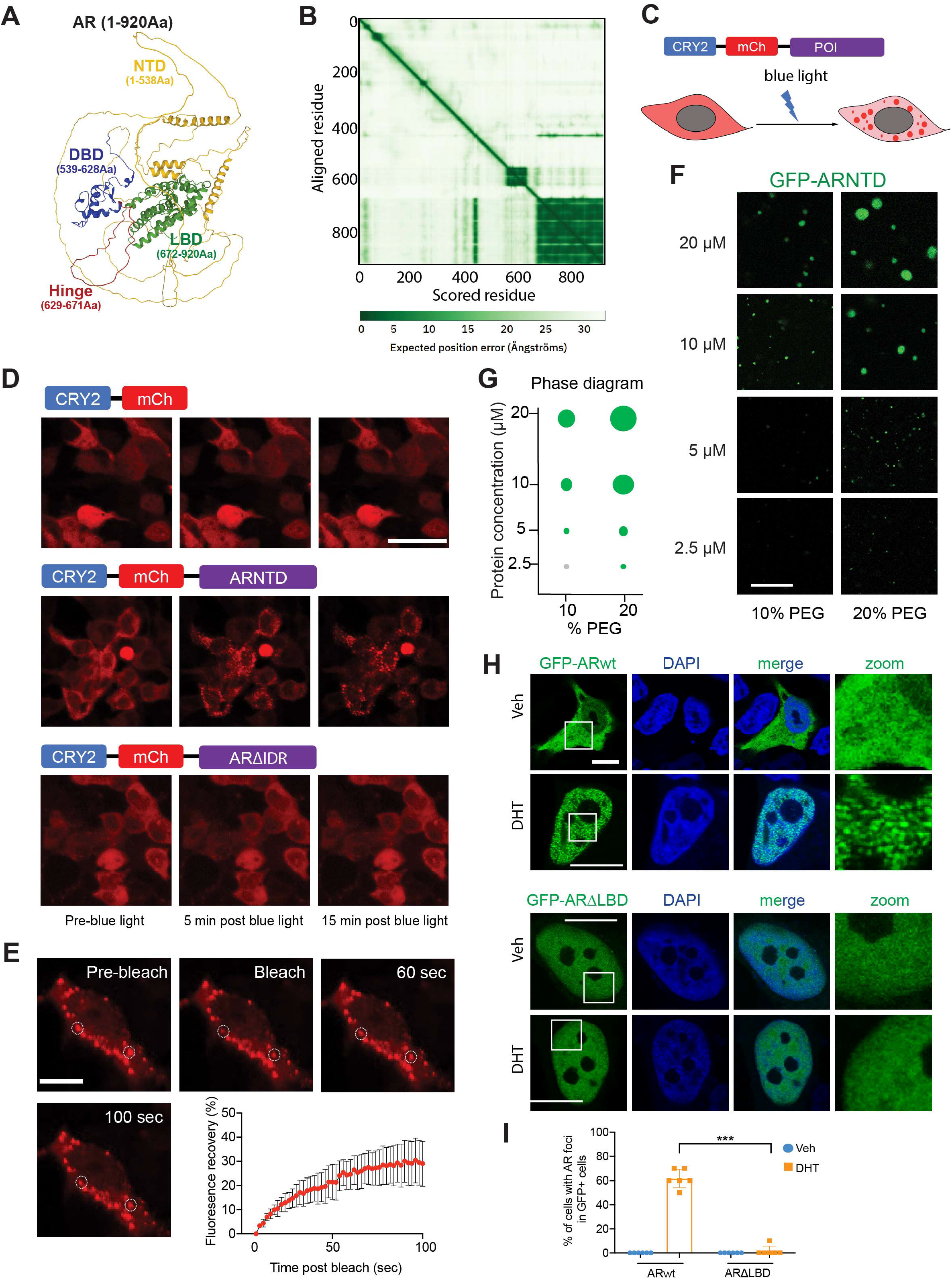
AR NTD is a low complexity domain and can drive phase separation in response to androgen signaling. **(A)** Structure prediction of full-length AR protein (1-920 Aa) by AlphaFold (https://alphafold.ebi.ac.uk/). Different domains of AR protein are indicated with matched font colors. **(B)** Predicted aligned error (PAE) plot indicating inter-domain accuracy. The shade of green shows expected distance error, with dark green indicating low error and light green indicating high error. **(C)** Schematic diagram of the optogenetic platform to study phase separation. The construct consists of the CRY2PHR domain, mCherry fluorescent protein and the protein of interest (POI). Upon 488nm blue light activation, CRY2 will rapidly cluster, resulting in condensation of proteins with phase separation capacity. **(D)** Representative images of 293T cells transiently transfected with CRY2-mCh control, CRY2-mCh-ARNTD, or CRY2-mCh-AR∆IDR constructs before and after blue light illumination. CRY2-mCh-ARNTD, but not CRY2-mCh-AR∆IDR, forms optoDroplets, suggesting phase separation capacity of ARNTD. Scale bar: 50 µm. **(E)** Representative images and fluorescence recovery curve of FRAP analysis on CRY2-mCh-ARNTD optoDroplets. Scale bar: 10 µm. **(F-G)** Representative droplet formation images and phase diagram of purified GFP-ARNTD at indicated protein and PEG8000 (PEG) concentrations. Scale bar: 10 µm. **(H-I)** Representative confocal images and quantification of LNCaP cells transiently transfected with GFP-ARwt or GFP-AR∆LBD and treated with vehicle or 100 nM DHT after cultured for 3 days in stripping medium. LBD deletion completely disrupted AR foci formation in response to DHT treatment. White boxes indicate the zoomed regions shown on the right. Scale bar: 10 µm. Percentages of cells showing AR foci in GFP-positive cells were plotted. Statistics: one-way ANOVA, ***P < 0.001.

We next examined AR behaviors in prostate cancer cell line LNCaP by transiently expressing GFP-tagged full-length AR (GFP-ARwt) in the cells. In the absence of DHT, GFP-ARwt was located predominantly in the cytoplasm with a relatively homogeneous distribution (Figure 1H). We then treated the transfected cells with 100 nM DHT and examined GFP-ARwt at various time points. After 10 min of DHT treatment, we started to see GFP-ARwt in the nucleus with a homogenous distribution. After 30 min of DHT treatment, GFP-ARwt was found in nuclei of all transfected cells, with about 30% of the cells having GFP-ARwt form numerous discrete assemblies. Here, we refer to the AR assemblies as condensates. Although the term condensate was originally derived from the phenomenon of phase separation, here we use its most recent definition - an entity that is not bound by a membrane, concentrating specific types of biomolecules, and involving non-stoichiometric, multivalent interactions (Banani et al., 2017; Choi et al., 2020; Mittag and Pappu, 2022) – without indicating phase separation as the underlying mechanism given the difficulty in rigorously proving phase separation in living cells (McSwiggen et al., 2019). The percentage of cells with GFP-ARwt foci reached to about 60% at 2 hours and did not further increase at 4 hours after DHT treatment (Figures 1H, 1I, and S1D). Therefore, we used 2-hour treatment for other in-cell AR condensates analyses.

Given that AR condensates formed in response to androgen stimulation, we tested if the ligand binding domain (LBD) of AR was required for AR condensation by transfecting a truncated AR lacking LBD (AR∆LBD) to LNCaP cells (Figure S1E). As there is a nuclear export sequence (NES) within the LBD (Saporita et al., 2003), GFP-AR∆LBD was located in the nucleus independent of DHT treatment. We found that GFP-AR∆LBD displayed a homogeneous distribution in both cells treated with vehicle control and cells treated with 100 nM DHT (Figures 1H and 1I). The requirement of LBD for AR to form condensates suggests that androgen stimulation through the structured LBD can induce multivalent interactions of AR NTD.

### AR requires its NTD to form condensates and to activate transcription

AR NTD is required for AR transactivation function and plays a critical role in the pathogenesis of prostate cancer, with 30% of AR mutations identified in prostate cancer mapped to the NTD (Gottlieb et al., 2012; Jenster et al., 1995; Monaghan and McEwan, 2016; Simental et al., 1991). To test if NTD is required for DHT-induced AR condensate formation, we expressed a GFP-tagged truncated AR protein without the N-terminal IDR (GFP-AR∆IDR). In the absence of DHT, GFP-AR∆IDR was in both cytoplasmic and nuclear compartments, with higher signal in the nucleus (Figures 2A and 2B). This was distinct from GFP-ARwt, which was predominantly located in the cytoplasm without DHT stimulation, suggesting that the NTD plays a role in retaining AR in the cytoplasm in the absence of androgen signal. In cells treated with DHT for 2 hours, GFP-AR∆IDR was located in the nuclei but did not form foci like full-length AR (Figures 2A, 2B, and 2C), indicating that AR requires its NTD to form condensates in response to androgen signal.

**Figure 2.**
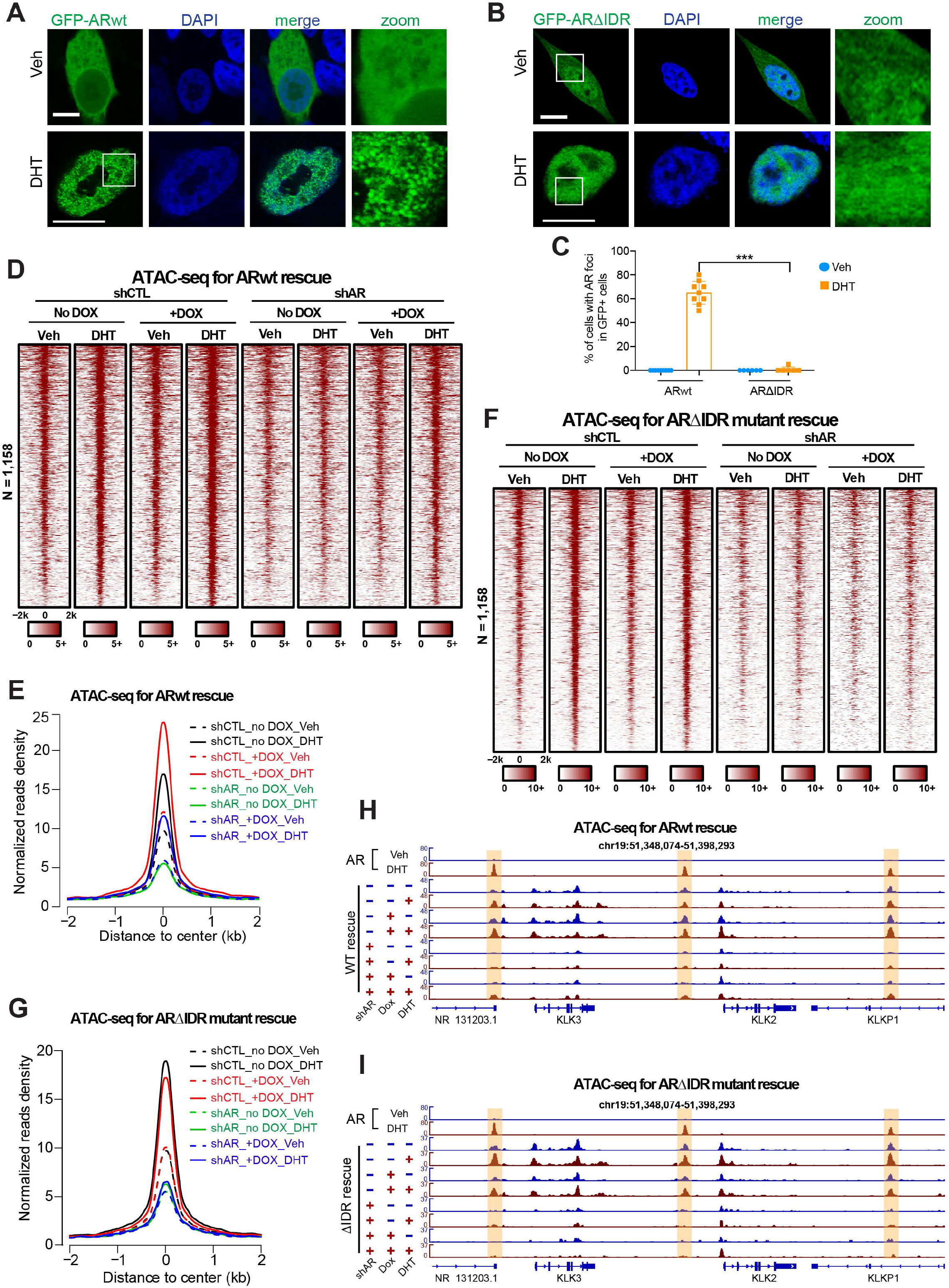
AR NTD is required for AR condensate formation and transcriptional activity. **(A-B)** Representative confocal images of LNCaP cells transiently transfected with GFP-ARwt (A) or GFP-AR∆IDR (B) treated with vehicle or 100 nM DHT. White boxes indicate the zoomed regions shown on the right. NTD deletion (∆IDR) abolished DHT-induced AR foci formation. Scale bar: 10 µm. **(C)** Quantification of LNCaP cells transiently transfected with GFP-ARWT or GFP-AR∆IDR with indicated treatments. Percentages of cells showing AR foci in GFP-positive cells were plotted. Statistics: one-way ANOVA, ***P < 0.001. **(D-E)** Heatmaps and aggregate plots of ATAC-seq data of LNCaP cells with indicated treatments to show chromatin accessibility of AR enhancers. LNCaP cells stably expressing Dox-inducible exogenous ARwt were transfected with control or AR shRNA to knock down endogenous AR. Exogenous ARwt expression was sufficient to promote chromatin opening and rescue the reduced chromatin accessibility caused by shAR. **(F-G)** Heatmaps and aggregate plots of ATAC-seq data of LNCaP cells stably expressing Dox-inducible exogenous AR∆IDR and transfected with control or AR shRNA to knock down endogenous AR. Exogenous AR∆IDR expression failed to promote chromatin opening and rescue the reduced chromatin accessibility caused by shAR. **(H)** Genome browser view of ATAC-seq signals on the enhancers of AR target genes KLK2 and KLK3 (highlighted by light yellow color). AR knockdown abolished DHT-induced chromatin opening and this phenotype was rescued by exogenous ARwt expression. **(I)** Genome browser view of ATAC-seq signals on the enhancers of AR target genes KLK2 and KLK3 (highlighted in light yellow) showing that IDR deletion abolished the enhancer activation function of AR.

An increasing amount of evidence has shown that IDR-IDR multivalent interactions that lead to TF hub or condensate formation is an important mechanism that drives transactivation (Boija et al., 2018; Cho et al., 2018; Chong et al., 2018; Hnisz et al., 2017; Nair et al., 2019; Sabari et al., 2018). We next looked for potential correlation between multivalent interactions and AR transcriptional activity by examining the requirement of NTD in AR-mediated enhancer activation and gene expression. To this end, we compared ARwt and AR∆NTD for their effects on chromatin accessibility of AR enhancers with ATAC-seq. 1,158 active AR enhancers were annotated using previously published AR ChIP-seq and H3K27ac ChIP-seq data (Hazelett et al., 2014; Malinen et al., 2017). To avoid any artifacts due to protein overexpression, we expressed AR under the control of a Tet-on promoter. As the endogenous AR protein might compete with exogenous AR and interfere with the comparison, we used shRNA to knockdown endogenous AR expression, followed by induction of exogenous AR expression which was resistant to shRNA due to synonymous mutations (Figures S2A, S2B, and S2C). As expected, in control shRNA groups, DHT stimulation significantly increased ATAC-seq signals on AR enhancers, and the expression of exogenous AR was sufficient to augment chromatin accessibility in both vehicle and DHT treated cells (Figures 2D and 2E). In shAR groups, DHT treatment had little effect on ATAC-seq signals, but exogenous AR expression was sufficient to restore DHT-induced chromatin opening to some level (Figures 2D and 2E), indicating that exogenous ARwt expression can rescue the defect caused by endogenous AR knockdown. In contrast, expression of AR∆IDR failed to increase ATAC-seq signals in control shRNA groups. Also, expression of AR∆IDR was unable to restore the diminished chromatin accessibility caused by shAR (Figures 2F and 2G), indicating that AR is dependent on its NTD to activate enhancers. As an example, the enhancer regions of the well-established AR target genes KLK2 and KLK3 were presented to show the effects of ARwt and AR∆IDR on enhancer activation (Figures 2H and 2I). The difference in the rescuing effects between ARwt and AR∆IDR was not due to the difference in expression levels, as we confirmed that their expression was comparable (Figures S2B and S2C).

To further evaluate the requirement of NTD for AR transcriptional activity, we performed RT-qPCR to measure AR target gene expression. Consistent with the ATAC-seq results, expressing ARwt in control shRNA groups significantly elevated the mRNA levels of KLK2 and NKX3-1. shAR abolished transcription of KLK2 and NKX3-1, but Dox-induced ARwt was sufficient to partially restore the transcription (Figure S2D). In contrast, expression of AR∆IDR has no effect on target gene transcription in the control shRNA group and failed to restore the reduced transcription caused by shAR (Figure S2E). Taken together, these data indicate that AR NTD is required for both condensate formation behaviors and transcriptional activity of AR.

### Disrupting the multivalent interactions of AR compromises AR-mediated transcription

The requirement of NTD (i.e., IDR) for both condensate formation and AR transcriptional activity prompted us to test if disrupting the multivalent IDR-IDR interactions inhibits AR-mediated transcription. We applied 1,6-Hexanediol (1,6-HD), an aliphatic alcohol that has been routinely used to disrupt multivalent hydrophobic protein-protein interactions to disassemble phase-separated droplets (Kato and McKnight, 2018; Lin et al., 2016), to LNCaP cells expressing GFP-ARwt. Treating the cells with 1,6-HD for 5 min after 2 hours of DHT treatment was sufficient to significantly reduce the percentage of cells exhibiting GFP-ARwt foci (Figures 3A and 3B). Similar results were found in U2OS human osteosarcoma cell line, which expresses AR and exhibits an androgenic response (Figures S3A and S3B). Like in LNCaP cells, GFP-ARwt distributed relatively homogeneously across the cytoplasm in vehicle control and formed discrete condensates in the nucleus upon DHT treatment in U2OS cells. The DHT-induced GFP-ARwt condensates disappeared when the cells were treated with 1,6-HD for 5 min (Figures S3A and S3B).

**Figure 3.**
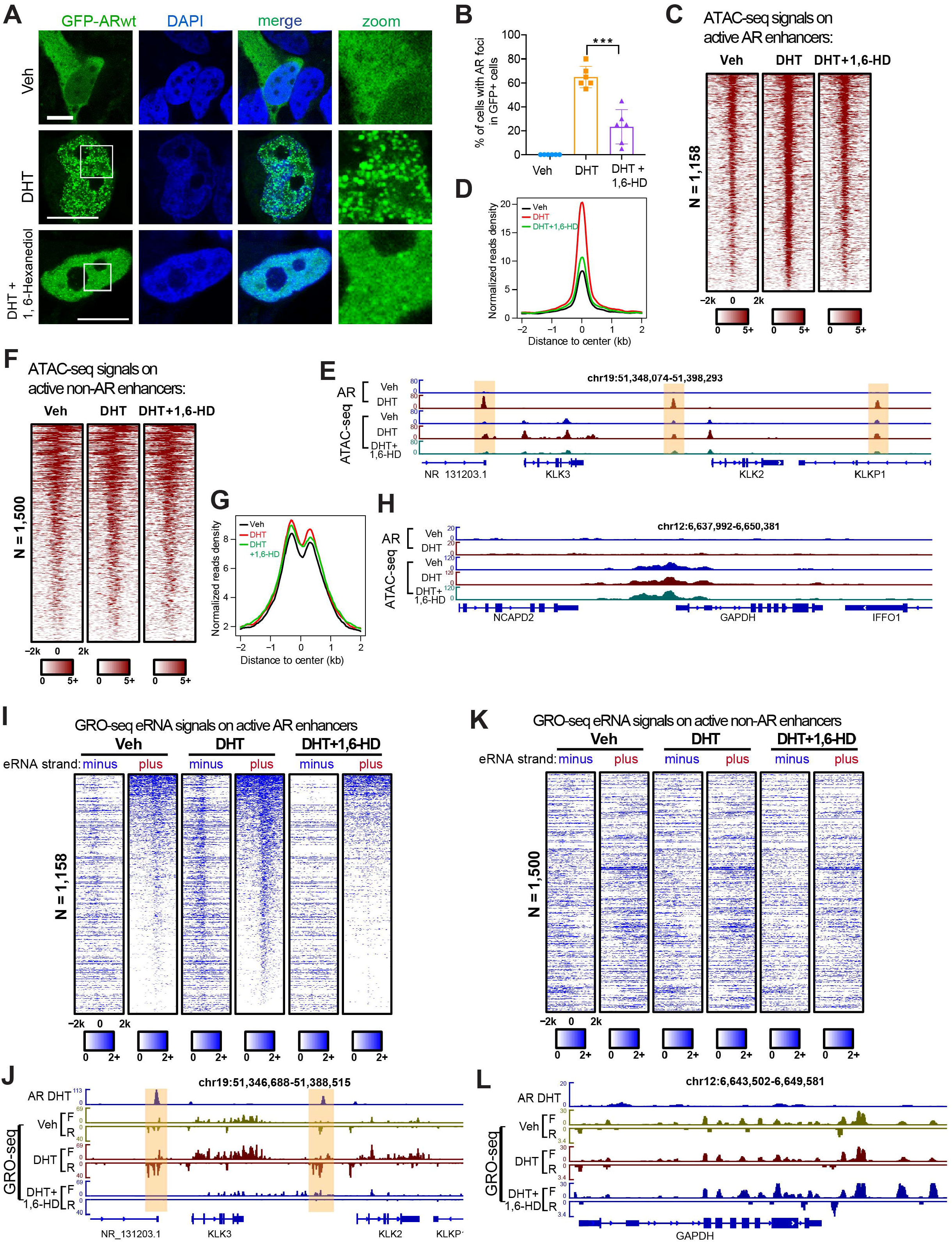
Disrupting the multivalent interactions of AR compromises AR-mediated transcription. **(A-B)** Representative confocal images and quantification of LNCaP cells transiently transfected with GFP-ARwt and treated with vehicle, DHT, or DHT+1,6-HD. 1,6-HD treatment significantly disrupted DHT-induced AR foci formation. White boxes indicate the zoomed regions shown on the right. Scale bar: 10 µm. Percentages of cells showing AR foci in GFP-positive cells were plotted. Statistics: one-way ANOVA, ***P < 0.001. **(C-D)** Heatmaps and aggregate plots of ATAC-seq signal on AR enhancers derived from LNCaP cells treated with vehicle, DHT, or DHT+1,6-HD. DHT treatment increased chromatin accessibility of AR enhancers and 1,6-HD inhibited chromatin opening in response to DHT. **(E)** Representative genome browser view of ATAC-seq signals on the enhancers of AR target genes KLK2 and KLK3 (highlighted in light yellow). **(F-G)** Heatmaps and aggregate plots of ATAC-seq signal on non-AR enhancers. DHT or 1,6-HD treatment did not affect the chromatin accessibility of non-AR enhancers. **(H)** Representative genome browser view of ATAC-seq signals on GAPDH gene locus as a negative control showing 1,6-HD did not affect the chromatin accessibility of a non-AR target gene. **(I-L)** Heatmaps and genome browser views of GRO-seq signals around the centers of AR active enhancer regions (I and J) or non-AR enhancer regions (K) or non-AR target gene GAPDH (L) under vehicle, DHT, or DHT+1,6-HD treatment conditions. DHT treatment promoted eRNA transcription on AR enhancers and this DHT effect was diminished by 1,6-HD treatment. AR enhancers were highlighted by light yellow color in (J).

We next examined the effect of 1,6-HD on chromatin accessibility of AR enhancers with ATAC-seq. We first confirmed that 1,6-HD treatment did not affect AR mRNA and protein levels in LNCaP cells (Figures S3C and S3D). We observed that DHT treatment dramatically increased the ATAC-seq signals on the 1,158 active AR enhancers (Figures 3C and 3D). This DHT-induced increase in chromatin accessibility was greatly abolished by 1,6-HD treatment. For instance, the enhancer regions of the AR target genes KLK2, KLK3 and NKX3-1 showed a low ATAC-seq signal in control cells and a high signal in DHT treated cells. Adding 1,6-HD to DHT treated cells reduced ATAC-seq signals on these regions (Figures 3E and S3E). As the negative controls, ATAC-seq signals on 1,500 active non-AR enhancers and GAPDH gene locus did not show obvious changes in response to DHT or 1,6-HD treatment (Figures 3F, 3G, and 3H). Therefore, these data suggest that disrupting the multivalent interactions of AR can compromise DHT-induced chromatin “opening” of AR enhancers.

We then performed GRO-seq (global run-on coupled with deep sequencing) to determine the impact of 1,6-HD on AR enhancer activity and target gene expression. Enhancer activity has been shown to highly correlate with enhancer RNA (eRNA) transcription, and eRNA production has been used as a reliable marker for enhancer activity (Hah et al., 2013; Lam et al., 2014). As expected, we detected increased eRNA synthesis from active AR enhancers upon DHT stimulation (Figure 3I), exemplified by KLK2, KLK3 and NKX3-1 enhancers (Figures 3J and S3F). Notably, 1,6-HD was sufficient to reduce DHT-induced eRNA transcription from active AR enhancers. As negative controls, transcription from active non-AR enhancers (Figure 3K) and GAPDH gene locus (Figure 3L) was not affected by 1,6-HD treatment. The inhibitory effects of 1,6-HD on AR enhancers activation and their associated AR target gene expression were also confirmed with RT-qPCR (Figure S3G), supporting the key role of AR multivalent interactions on regulating enhancer activity and target gene transcription.

### Multivalent homotypic interactions and transcriptional activity of AR depend on the aromatic residues within its NTD

Recent studies have shown that aromatic residues in IDRs can function as “stickers” to promote multivalent interactions that underlie phase separation (Ho and Huang, 2022; Kwon et al., 2013; Li et al., 2018; Lin et al., 2017; Wang et al., 2018). There are seven phenylalanine residues within AR NTD. To evaluate their contribution to the multivalent interaction and phase separation behaviors of AR, we substituted all seven phenylalanine residues with serine amino acids (AR7FS) (Figure S4A). We first expressed GFP-AR7FS in LNCaP cells and examined its distribution. Like GFP-ARwt, GFP-AR7FS was homogeneously distributed in the cytoplasm in the absence of DHT and was detected exclusively in nuclei upon DHT treatment. However, the DHT-induced condensate formation was significantly disrupted by the 7FS mutation (Figures 4A and 4B). We next performed the *in vitro* droplet formation assay for recombinant GFP-NTD7FS fragment. Although GFP-NTD7FS formed droplets in the presence of the crowding agent PEG8000, the droplet size was significantly smaller compared to that of GFP-NTDwt droplets at all protein concentrations tested (Figures 4C and 4D). Together, these results indicate that the phenylalanine residues play a key role in the multivalent homotypic interactions of AR.

**Figure 4.**
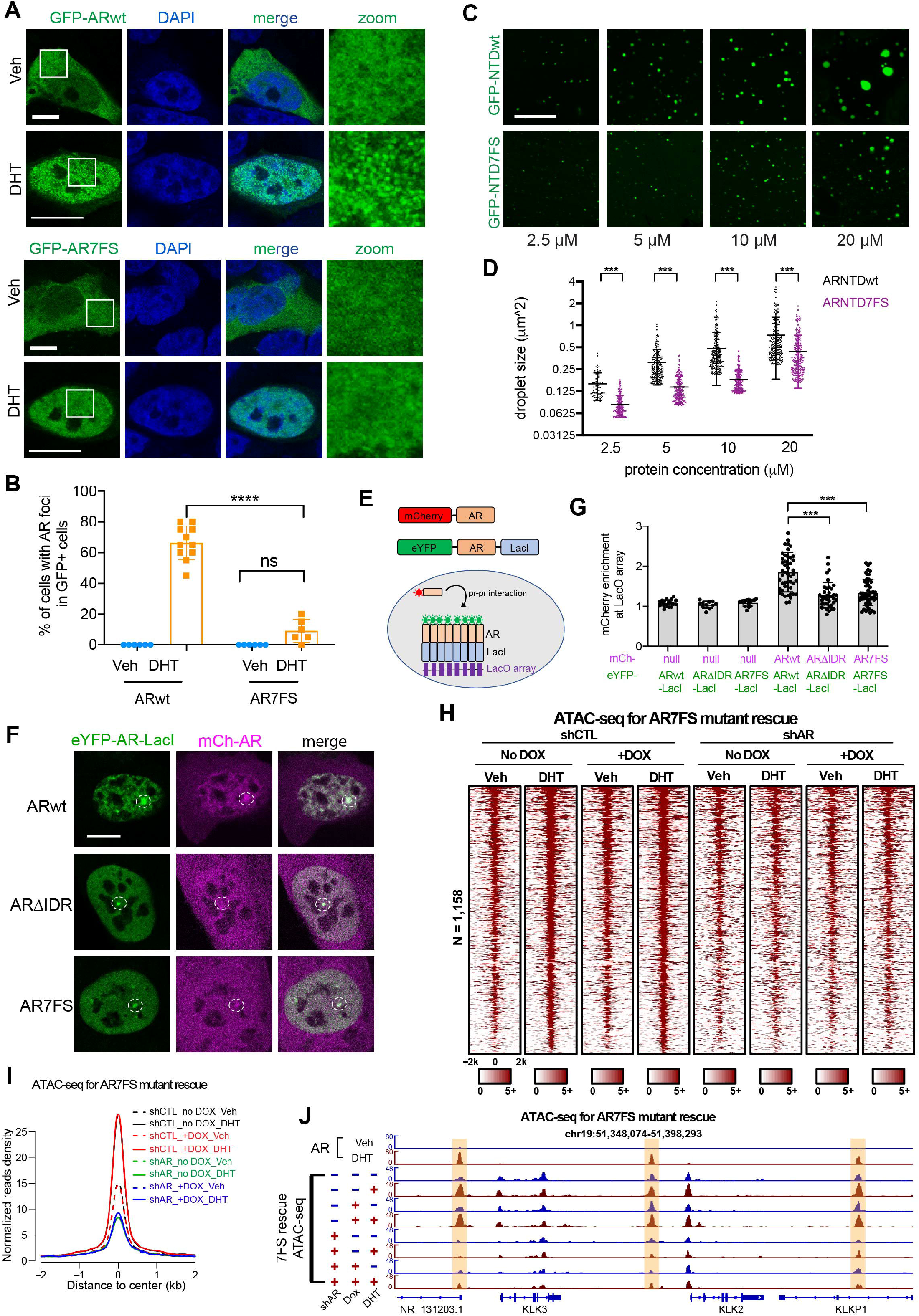
Aromatic residues mutation in NTD weakens multivalent AR-AR interactions and disrupts AR transcriptional activity. **(A-B)** Representative confocal images and quantification of LNCaP cells transiently transfected with GFP-ARwt or GFP-AR7FS and treated with vehicle or DHT. Mutating the aromatic residues inhibited DHT-induced AR condensate formation. White boxes indicate the zoomed regions shown on the right. Scale bar: 10 µm. Percentages of cells showing AR foci in GFP-positive cells were plotted. Statistics: one-way ANOVA, ***P < 0.001. **(C)** Representative droplet formation images of purified GFP-NTDwt and GFP-NTD7FS at indicated protein concentrations. Scale bar: 10 µm. **(D)** Quantification of the size of droplets formed by purified GFP-NTDwt and GFP-NTD7FS at indicated protein concentrations. Statistics: one-way ANOVA, ns: non-significant, ****P < 0.0001. **(E)** Schematic illustration of the LacO array system to test AR self-interactions. eYFP-AR-LacI is recruited to LacO array through protein-DNA binding and mCherry-AR can be recruited to the LacO array through AR-AR interactions. **(F)** Representative images of LacO array-containing U2OS cells co-expressing indicated proteins. Scale bar: 10 µm. **(G)** Quantification of mCherry-AR recruitment to the LacO hub through AR-AR self-association. Enrichment of mCherry above relative level of 1 suggests AR-AR self-association. Statistics: one-way ANOVA, ***P < 0.001. **(H-I)** Heatmaps and aggregate plots of ATAC-seq signal on AR enhancers derived from LNCaP cells with indicated treatments. Exogenous expression of AR7FS failed to rescue the reduced chromatin accessibility on AR enhancers caused by shAR. **(J)** Genome browser view of ATAC-seq signals on the enhancers of AR target genes KLK2 and KLK3 (highlighted in light yellow) showing that 7FS mutation abolished the enhancer activation function of AR.

Given the correlation between the multivalent homotypic interactions of AR and its transcriptional activity, we sought to test if such interactions promote its recruitment to chromatin using our previously established cell imaging assay (Chong et al., 2018; Chong et al., 2022; Wan et al., 2020). We co-expressed mCherry-AR and eYFP-AR-LacI in U2OS 2-6-3 cells containing a synthetic Lac operator (LacO) array integrated into the genome (Janicki et al., 2004). Through targeted DNA binding, eYFP-ARwt-LacI molecules were recruited to the LacO array, generating a large concentrated interaction hub on the chromatin (Figures 4E and 4F). We observed a strong mCherry signal at the array (Figures 4F and 4G), indicating a strong homotypic interaction ability of ARwt. When we performed the LacO assay using AR∆IDR, we observed a much weaker mCherry signal at the array, even though the recruitment of eYFP-AR∆IDR-LacI molecules to LacO array appeared normal (Figures 4F and 4G), consistent with the essential role of IDR in mediating AR homotypic interactions. Notably, the 7FS mutation also significantly reduced AR homotypic interaction to a level similar to AR∆IDR (Figures 4F and 4G). Collectively, these data suggest that aromatic residues in the NTD promote the enrichment of AR to chromatin through its multivalent homotypic interactions.

We next asked how 7FS mutation affected DHT-induced enhancer activation with ATAC-seq. We knocked down endogenous AR expression and expressed exogenous AR7FS (Figure S4B) to see if AR7FS could rescue the defects in chromatin accessibility caused by shAR. Consistent with data above (Figures 2D and 2F), shAR abolished DHT-induced chromatin opening on AR enhancers. Unlike ARwt which was sufficient to restore the ATAC-seq signals (Figures 2D, 2E, and 2H), AR7FS was unable to rescue the ATAC-seq phenotype caused by shAR (Figures 4H, 4I, and 4J). Similarly, shAR disrupted AR target gene transcription and expression of AR7FS failed to rescue the defects (Figure S4C). Therefore, the aromatic residues within the NTD are critical for AR multivalent interactions and transcriptional activity.

### AR NTD can be functionally substituted by FUS and TAF15 IDRs, but not ERα IDR

Previous studies have shown that in some cases one IDR can functionally substitute for another in RNP granule assembly (Decker et al., 2007; Gilks et al., 2004). We wondered whether other IDRs could substitute for AR NTD to regulate AR condensate formation and transcriptional activity. Fused in sarcoma/translocated in liposarcoma (FUS/TLS or FUS) and TAF15 represent two of the most studied examples of proteins that undergo multivalent interactions and phase separation (Portz et al., 2021). The prion-like domain (PrLD) in FUS protein is an IDR enriched with tyrosine residues and can undergo phase separation driven by tyrosine-tyrosine interactions (Wang et al., 2018). TAF15 IDR has the same number of tyrosine residues but more charged residues than FUS IDR and exhibits a strong tendency to phase separate (Wei et al., 2020). We replaced AR NTD with the IDR from FUS or TAF15 and examined AR condensate formation in LNCaP cells (Figure S5A). As mentioned above, GFP-ARwt showed a homogeneous distribution in the cytoplasm in vehicle control but localized in the nucleus and became punctate upon DHT treatment. NTD deletion promoted nuclear localization in the absence of DHT and abolished the DHT-induced condensate formation (Figures 5A and 5B). Fusion of FUS IDR or TAF15 IDR to AR∆IDR (GFP-FUSIDR-AR∆IDR or GFP-TAF15IDR-AR∆IDR) fully restored DHT-induced AR condensate formation. Notably, FUS IDR and TAF15 IDR were able to promote AR condensate formation in some cells even in the absence of DHT (Figures 5A and 5B). We further tested if IDR from ERα, another hormone-related transcription factor like AR, could substitute for AR IDR (Figure S5A). ERα has been previously shown to undergo phase separation and to promote enhanceosome assembly (Nair et al., 2019). Fusion of ERα IDR to AR∆IDR (GFP-ERαIDR-AR∆IDR) did not affect AR∆IDR distribution in the vehicle condition, but partially restored DHT-induced condensate formation. We observed a similar percentage of cells showing puncta in cells expressing GFP-ERαIDR-AR∆IDR after DHT treatment (Figures 5A and 5B). However, these condensates were less distinct from background compared to GFP-ARwt condensates (Figure 5A). This was also reflected by the fringe visibility measurement (Figure 5B).

**Figure 5.**
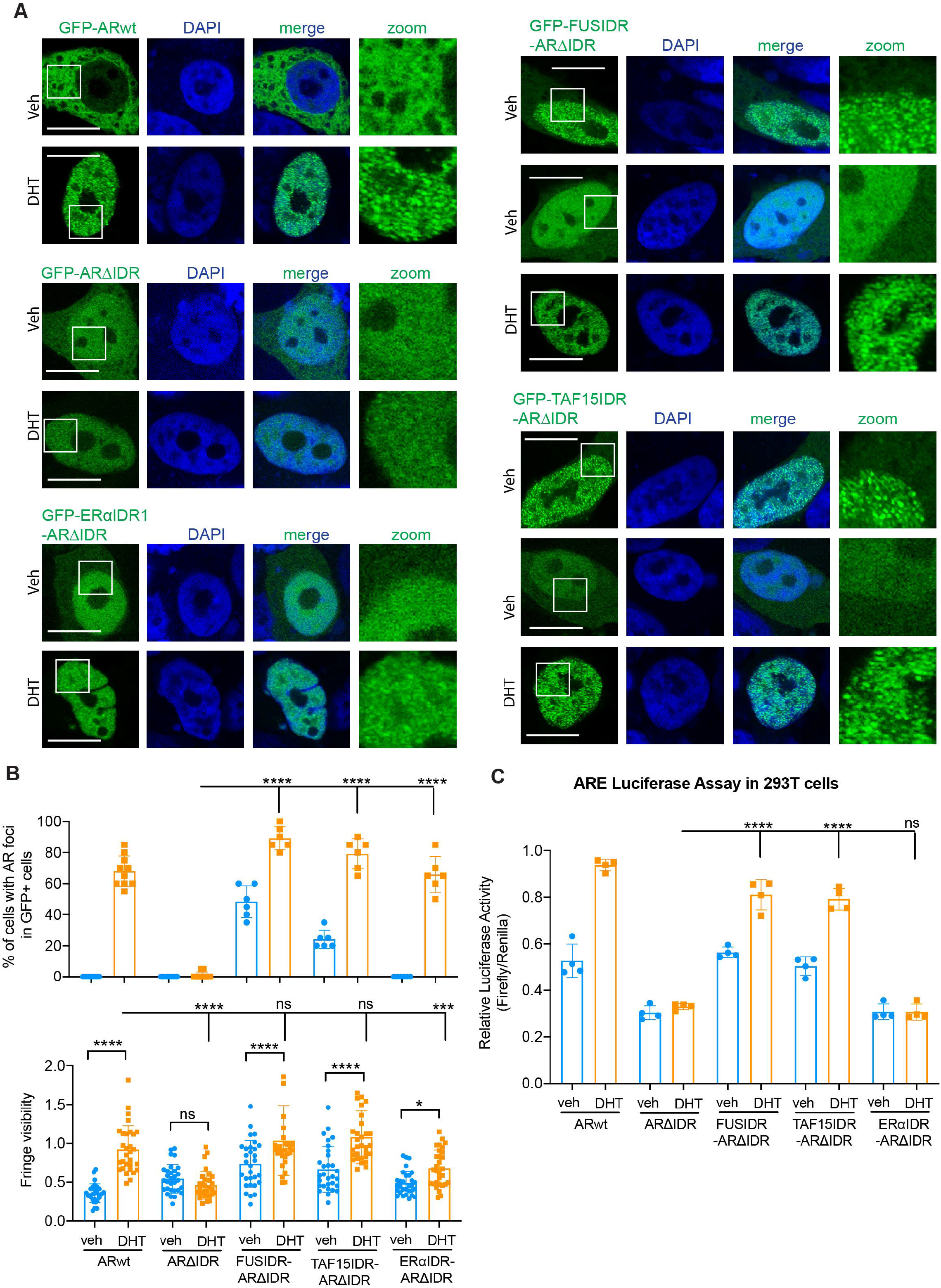
AR NTD can be functionally substituted by FUS and TAF15 IDRs, but not ERα IDR. **(A)** Representative confocal images of LNCaP cells transiently transfected with the indicated AR constructs and treated with vehicle or DHT. Replacing ARIDR with FUSIDR or TAF15IDR retained the DHT-induced condensate formation capacity of AR. FUSIDR and TAF15IDR also promoted AR condensate formation in some cells even in the absence of DHT. White boxes indicate the zoomed regions shown on the right. Scale bar: 10 µm. **(B)** Quantifications of LNCaP cells with AR condensates (top) and the fringe visibility of AR condensates (bottom). In the top panel, percentages of cells showing AR foci in GFP-positive cells were plotted. In the bottom panel, fringe visibility values of randomly selected AR foci from 6 cells (5 foci/cell) were plotted for each condition. Statistics: one-way ANOVA, ns: non-significant, ****P < 0.0001, ***P < 0.001, *P < 0.05. **(C)** Luciferase reporter assay to examine transcriptional activity of the indicated AR proteins and conditions. 293T cells were co-transfected with the luciferase reporter vector containing 3xARE and one of the AR constructs followed by luciferase activity measurement. FUSIDR and TAF15IDR, but not ERαIDR, were able to rescue the disrupted transcriptional activity caused by AR IDR deletion. Statistics: one-way ANOVA, ns: non-significant, ****P < 0.0001.

As AR NTD can be substituted by other IDRs for its role in promoting condensate formation, we next tested if AR NTD can be replaced for its transcriptional activity. For this purpose, we co-expressed a luciferase reporter containing 3xARE with various AR proteins to examine their transcriptional activity. As expected, expression of ARwt significantly increased luciferase activity in response to DHT treatment, indicating the strong transcriptional activity of ARwt (Figure 5C). Consistent with its impaired transcriptional activity (Figure 2), AR∆IDR failed to promote luciferase gene expression upon DHT stimulation. Remarkably, the fusion proteins FUSIDR-AR∆IDR and TAF15IDR-AR∆IDR were both as efficient as ARwt in promoting luciferase expression in response to DHT. In contrast, ERαIDR-AR∆IDR was insufficient to activate luciferase expression (Figure 5C).

We further tested the fusion proteins for their function in regulating AR target gene transcription in LNCaP cells with endogenous AR knocked down (Figures S5B, S5C and S5D). RT-qPCR data showed that expression of exogenous FUSIDR-AR∆IDR and TAF15IDR-AR∆IDR was sufficient to promote DHT-induced target gene expression after shAR knockdown (Figures S5E and S5F). Consistent with the results from luciferase reporter assay (Figure 5C), ERαIDR-AR∆IDR behaved like AR∆IDR and was unable to promote target gene expression (Figures S2E and S5G). Together, these results indicate that AR IDR can be functionally substituted by selective IDRs. The correlation of condensate formation status and transcriptional activities of these fusion proteins also support that the biophysical properties of AR condensates are critical for its transcriptional activity.

### PolyQ track expansion leads to more stable AR condensates and reduces AR transcriptional activity

PolyQ expansion has been recently shown to promote condensate formation. For instance, the polyQ tract of huntingtin protein can drive reversible liquid-like assemblies, which can be then converted to solid-like assemblies with increased polyQ length (Peskett et al., 2018). There is a polymorphic number of glutamine (Q) repeats in AR NTD and the length of polyQ tract is believed to contribute to individual differences in androgen sensitivity, with shorter polyQ tracts associated with higher AR transcriptional activity (Monks et al., 2007). Q repeats beyond the normal range (>36Q) is associated with Kennedy’s disease/spinal and bulbar muscular atrophy (KD/SBMA). We thus used AR with 69 Q repeats (ARpQ69) to study the effects of polyQ expansion on AR condensate formation and transcriptional activity. We first performed the *in vitro* droplet formation assay to compare ARNTDpQ69 to ARNTDwt that contains 23 Q repeats. Similar to GFP-NTDwt, GFP-NTDpQ69 was able to form droplets in the presence of PEG8000 and the droplet size correlated with protein concentration (Figures 6A and 6B). Notably, GFP-NTDpQ69 formed slightly larger droplets compared to GFP-NTDwt at all protein concentrations tested. And the saturation concentration (the lowest concentration at which droplets appear) for GFP-NTDpQ69 was lower than that of GFP-NTDwt (Figures 6A and 6B). At 20 uM protein concentration, some GFP-NTDpQ69 droplets appeared to aggregate into large assemblies with irregular shapes (Figure 6A). Together, these results suggest that polyQ expansion might enhance the phase separation propensity of AR.

**Figure 6.**
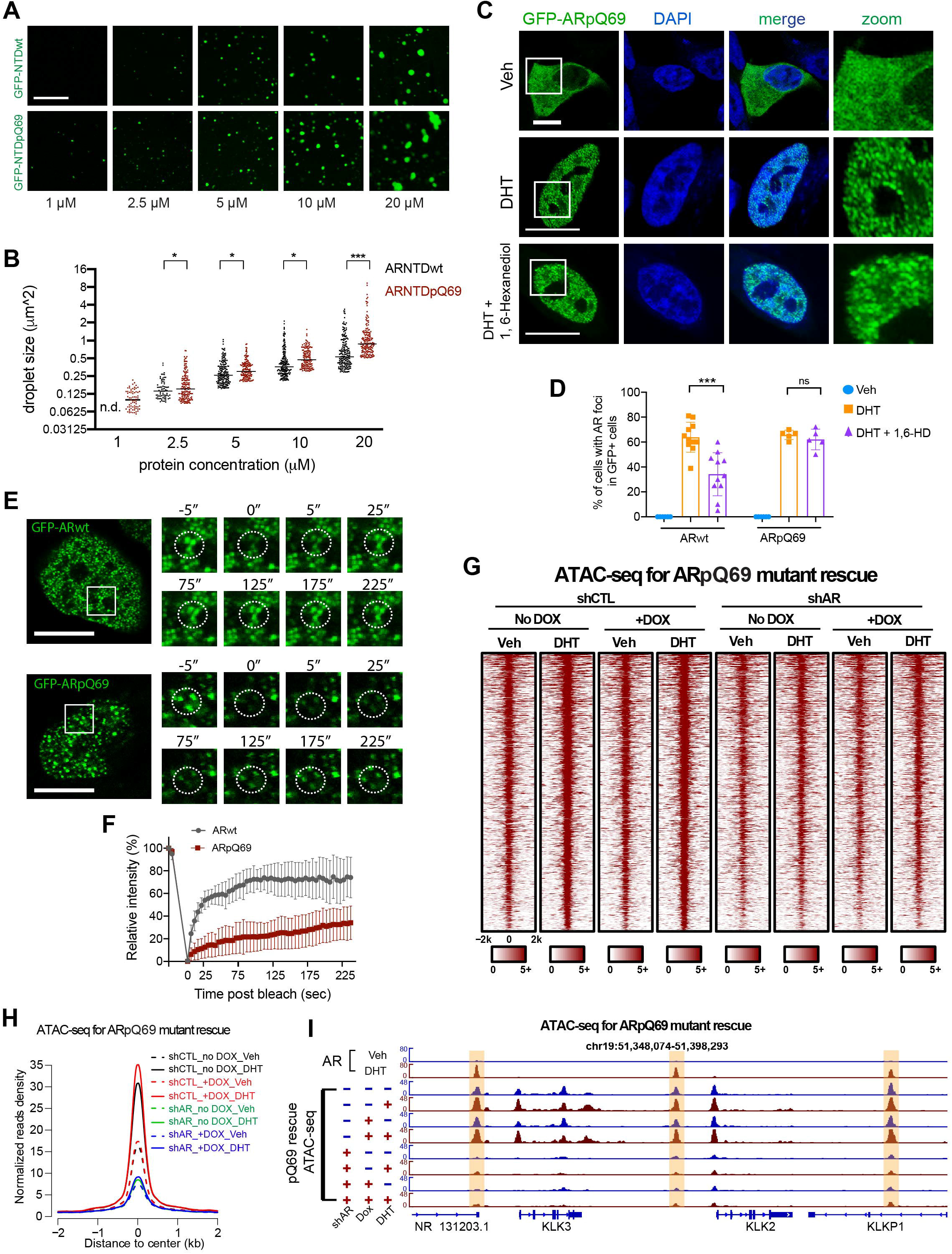
PolyQ expansion leads to more stable AR condensates and reduces AR transcriptional activity. **(A)** Representative droplet formation images of purified GFP-NTDwt (pQ23) and GFP-NTDpQ69 at indicated protein concentrations. Scale bar: 10 µm. **(B)** Quantification of the size of droplets formed by purified GFP-NTDwt and GFP-NTDpQ69 at indicated protein concentrations. n.d.: non-detectable. Statistics: student’s t-test, *P < 0.05, ***P < 0.001. **(C-D)** Representative confocal images and quantification of LNCaP cells transiently transfected with GFP-ARpQ69 and treated with vehicle, DHT, or DHT+1,6-HD. Scale bar: 10 µm. Percentages of cells showing AR foci in GFP-positive cells were plotted. GFP-ARpQ69 foci were more resistant to 1,6-HD treatment compared to GFP-ARwt. Statistics: one-way ANOVA, ns: non-significant, ***P < 0.001. **(E-F)** Representative images and quantification of Fluorescence Recovery After Photobleaching (FRAP) analyses on GFP-ARwt and GFP-ARpQ69 condensates formed in LNCaP cells in response to DHT treatment. Scale bar: 10 µm. Error bar: standard deviation. **(G-H)** Heatmaps and aggregate plots of ATAC-seq signals around the centers of AR active enhancer regions under the indicated conditions. ARpQ69 was not able to substitute for ARwt for AR function in activating AR enhancers. **(I)** Genome browser view of ATAC-seq signals on the enhancers of AR target genes KLK2 and KLK3 (highlighted by light yellow color) showing that polyQ track expansion abolished the enhancer activation function of AR.

We next tested if polyQ expansion can affect AR condensate formation in cells. We observed that, like GFP-ARwt, GFP-ARpQ69 underwent DHT-induced nuclear localization and condensate formation. Interestingly, GFP-ARpQ69 condensates were more resistant to 1,6-Hexanediol treatment than GFP-ARwt (Figures 3A, 6C, and 6D), indicating differences in the chemistry underlying the multivalent interactions of ARwt and ARpQ69. When we performed FRAP analyses, we found that GFP-ARpQ69 condensates had a slower and far less complete recovery after photobleaching than those of GFP-ARwt (Figures 6E and 6F), suggesting more stable multivalent protein-protein interactions of ARpQ69.

We then assessed the effect of polyQ expansion on AR function in enhancer activation with ATAC-seq. We expressed Dox-inducible exogenous ARpQ69 in LNCaP cells with endogenous AR knocked down (Figures S6A and S6B). While exogenous ARwt expression was sufficient to rescue the defects in chromatin accessibility caused by AR knockdown (Figures 2D, 2E, and 2H), ARpQ69 expression failed to replace endogenous AR to mediate DHT-induced chromatin opening on AR enhancers (Figures 6G, 6H, and 6I). Consistent with the ATAC-seq result, our RT-qPCR data showed that ARpQ69 was insufficient to promote AR target gene transcription (Figure S6C). Using the luciferase reporter with 3xARE, we found that ARpQ69 was unable to activate luciferase expression, just like AR∆IDR and AR7FS (Figure S6D). Therefore, despite its strong condensate formation capacity, ARpQ69 exhibits impaired transcriptional activity. These results support the notion that an optimal level of AR-AR multivalent interactions is required for its proper function, consistent with recent findings on other endogenous and synthetic TFs (Chong et al., 2022; Trojanowski et al., 2022).

### An optimal level of AR multivalent interactions is critical for the assembly of AR enhancer complex

The observations above indicate that the biophysical properties of AR condensates influence AR function. To understand the underlying mechanisms, we first tested whether the IDR-mediated condensate formation plays a role in AR binding on AR enhancers. We expressed AR in LNCaP cells with endogenous AR knocked down and performed ChIP-seq using the BirA-BLRP system (Liu et al., 2014) to profile AR occupancy on chromatin (Figure S7A). As expected, ARwt exhibited strong binding on active AR enhancers (Figures 7A and 7B). Interestingly, we observed a much weaker binding strength of AR∆IDR than ARwt (Figures 7A and 7B).

**Figure 7.**
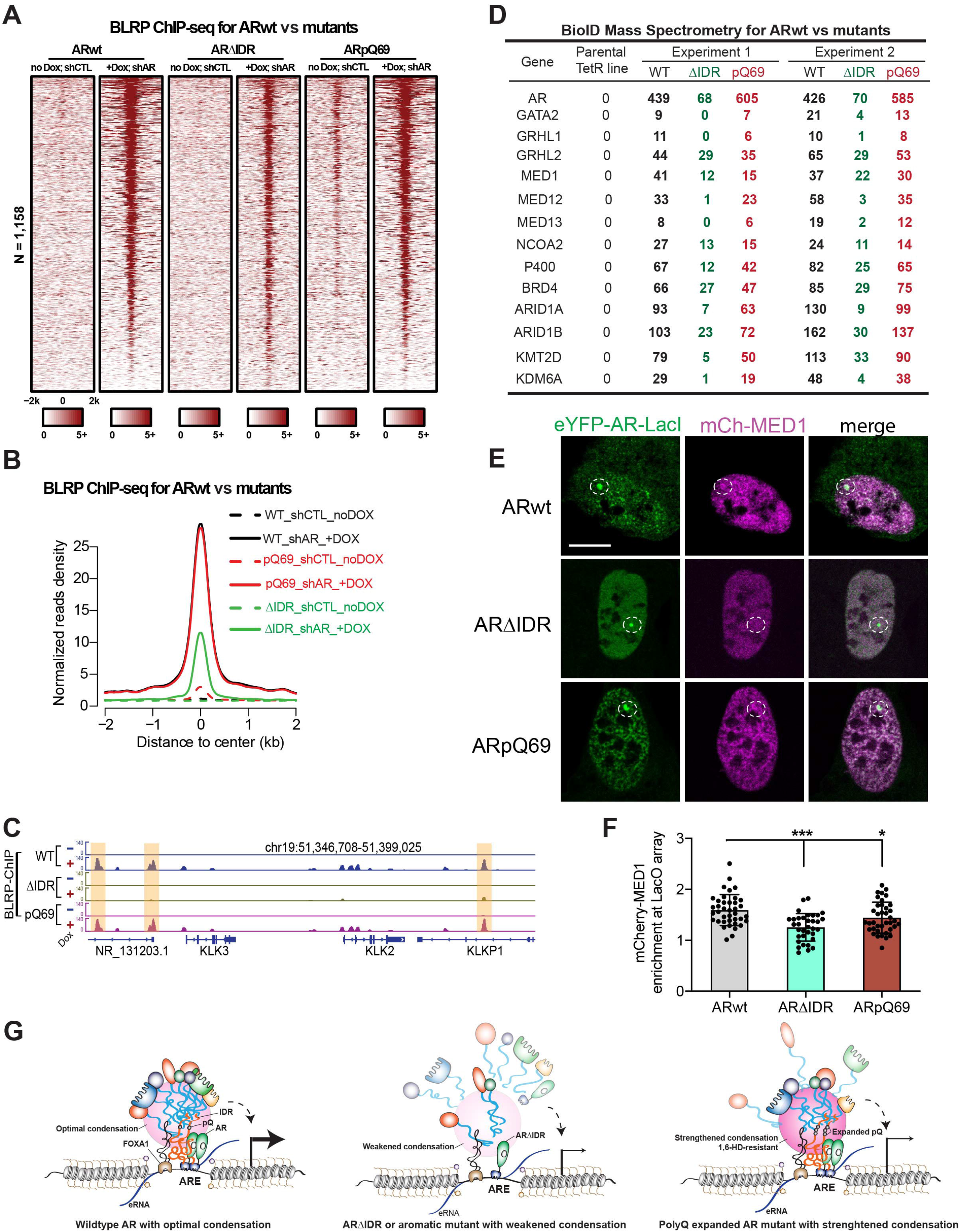
An optimal level of AR multivalent interactions is critical for the assembly of AR enhancer complex. **(A-B)** Heatmaps and aggregate plots of AR ChIP-seq data showing the binding of ARwt, AR∆IDR and ARpQ69 on active AR enhancers. ChIP-seqs were performed using BirA-BLRP LNCaP stable cell lines in which endogenous AR was knocked down with AR shRNA and exogenous expression of BLRP-tagged AR proteins was induced by doxycycline. **(C)** Genome browser view of ChIP-seq signals on the enhancers of AR target genes KLK2 and KLK3 (highlighted by light yellow color) showing that IDR deletion abolished AR binding but polyQ track expansion did not affect AR binding on AR enhancers. **(D)** A list of enhancer complex components identified from our BioID analyses using ARwt, AR∆IDR or ARpQ69 as a bait protein. Normalized peptide numbers of each identified protein were listed for each experiment. IDR deletion and polyQ expansion both reduced the peptide number of enhancer component proteins in AR complex. **(E)** Representative images of LacO array-containing U2OS cells co-expressing indicated proteins. Scale bar: 10 µm. **(F)** Quantification of mCherry-MED1 recruitment to the LacO hub through AR-MED1 association. Enrichment of mCherry above relative level of 1 suggests AR-MED1 heterotypic interactions. Statistics: one-way ANOVA, ***P < 0.001, *P < 0.05. **(G)** Schematic working model for AR enhancer assembly. An optimal level of AR-AR or AR-cofactor multivalent interactions mediated by wildtype AR IDR promotes the assembly of enhancer complex to activate transcription. Disturbing (e.g., ∆IDR or 7FS mutation) or overly-strengthening AR condensation (e.g., pQ69) leads to defective enhancer assembly and disrupted transcriptional activation.

As AR∆IDR contained an intact DBD, the reduced binding of AR∆IDR suggested that IDR-mediated AR condensate formation can facilitate AR-enhancer interaction. This is consistent with a recent report that IDRs guide TF binding on chromatin by localizing TFs to broad DNA regions surrounding the binding sites (Brodsky et al., 2020). Notably, ARpQ69 showed similar binding capacity onto AR enhancers as ARwt (Figures 7A and 7B), exemplified by their binding patterns on KLK2, KLK3, and NKX3-1 enhancers (Figures 7C and S7B). As ARpQ69 showed normal chromatin occupancy but disrupted AR function (Figure 6), we asked whether ARpQ69 had defects in enhancer assembly. Through the well-established BioID approach (Bi et al., 2020; Roux et al., 2012; Zhu et al., 2019) with ARwt as the bait protein, we identified a series of enhancer complex components as AR-interacting proteins, including TFs, epigenetic cofactors, and chromatin remodelers (Figures S7C and 7D). When we performed BioID using AR∆IDR as a bait, we detected a lower peptide number of AR∆IDR bait due to the shorter AR protein length and identified the same set of enhancer complex components. However, the peptide numbers of these AR cofactors were dramatically lower compared to ARwt BioID, indicating that IDR is required for AR enhancer complex assembly (Figures S7C and 7D). ARpQ69 BioID also recognized the same set of AR-interacting enhancer components, and their peptide numbers were significantly lower than those in ARwt BioID despite the higher peptide number of the bait protein (Figures S7C and 7D). We next performed Gene Ontology enrichment analyses on AR-interacting proteins that showed reduced interaction with AR∆IDR or ARpQ69 compared to ARwt and identified the enrichment of proteins involved in mRNA processing and transcription (Figures S7D and S7E). Therefore, even though polyQ tract expansion does not affect AR binding to enhancers, it might interfere with AR enhancer assembly, leading to reduced transcriptional activity. To further test this possibility, we measured heterotypic interactions between AR and MED1, a transcriptional coactivator in the enhancer complex, using the LacO array system. We co-expressed mCherry-MED1 and eYFP-ARwt-LacI in U2OS cells containing the LacO array and observed strong mCherry enrichment at the array (Figures 7E and 7F), indicating AR-MED1 interaction. When we performed the assay with AR∆IDR or ARpQ69, we observed significantly reduced enrichment of mCherry-MED1 (Figures 7E and 7F), suggesting that deletion of AR IDR or expansion of polyQ tract impaired the heterotypic AR-MED1 interaction. These data collectively support a model that AR needs to undergo multivalent homotypic interactions at an optimal level to enable its recruitment of enhancer components to activate enhancers, and that weakening or overly strengthening AR condensation impairs enhancer assembly, resulting in reduced transcription of AR target genes (Figure 7G).

## Discussion

AR responds to androgen signaling and enters the nucleus to assemble active enhancers that consists of various factors including epigenetic co-activators and other DNA-binding TFs. However, it is not clear how AR interacts with other factors to assemble active enhancers. Here, we report that AR forms local high-concentration condensates driven by its IDR-mediated multivalent interactions in response to androgen stimulation and that disturbing AR condensates impairs AR function in enhancer assembly. More surprisingly, we find that extending the polyQ repeats within AR IDR can strengthen AR-AR multivalent interactions, also resulting in impaired AR function in enhancer assembly. Our data reveal that AR enhancer assembly requires an optimal level of AR multivalent interactions to mediate cofactor recruitment, and that altering the biophysical properties of AR condensates changes the involved homotypic and heterotypic interactions.

Most enhancer complex proteins, including TFs and their cofactors, comprise both structured domains and IDRs. At the chromatin level, the contributions of structured domains on protein-DNA and protein-protein interactions have been well studied. However, our investigation on the roles of IDRs of enhancer complex proteins is relatively recent. Despite the emerging evidence that convincingly supports the important role of IDR-mediated multivalent interactions in regulating enhancer activation, many questions remain unanswered. How do so many different IDRs interact to activate specific enhancers? How do those structured domains interact with IDRs to modulate protein behaviors? How do different signals impact IDR interactions and protein functions? Our work on DHT-induced AR multivalent interactions through its IDR provides important insights into these questions.

The requirement of structured LBD for AR condensate formation (Figure 1H) indicates that the structured domains may contribute to hormone-induced multivalent interactions of IDR. AR LBD contains 11 α-helices that form the androgen-binding pocket (Matias et al., 2000). Upon androgen binding, the LBD changes its conformation to become an activation function domain (AF-2) to recruit coregulators with LXXLL motifs (Heery et al., 2001). The ^23^FQNLF^27^ sequence within AR NTD can act similarly as an LXXLL motif (He et al., 2004). Ligand binding induces an intramolecular N/C interaction between LBD and ^23^FQNLF^27^, followed by nuclear translocation. Inside the nucleus, AR dimerizes through the intermolecular N/C interaction (van Royen et al., 2007). It’s thus likely that intramolecular interaction between AR NTD and the androgen-bound AR LBD can promote AR intermolecular multivalent interactions and condensate formation. Consistent with this notion, it has been reported that estrogen binds to ERα LBD to promote ERα co-separation with MED1 (Boija et al., 2018). Therefore, we propose that the binding of hormone agonists or hormone antagonists at structured LBD may alter IDR behaviors of hormone receptors. Several different androgen antagonists have been developed to target LBD, including bicalutamide, flutamide, nilutamide and enzalutamide. It would be interesting to test if these antagonists regulate the condensate formation behavior of AR and affect AR enhancer assembly through influencing IDR-IDR or IDR-structured domain interactions.

Another interesting observation in the study is that AR IDR can be functionally substituted by FUS and TAF15 IDRs, but not ERαIDR (Figure 5). Chimeric proteins with FUS/TAF15 IDR and AR∆IDR formed condensates in LNCaP cells upon DHT treatment. They displayed the same level of transcriptional activity as wildtype AR in the luciferase reporter assay. Although the chimeric protein with ERαIDR and AR∆IDR also formed condensates in cells, the condensates were less distinct than the ones of ARwt, as measured by the fringe visibility, and it showed no transcriptional activity. One possible reason for the difference between FUS/TAF15 IDRs and ERαIDR might be their distinct multivalent interaction capacity and different signaling dependency. ERαIDR might require the interaction with estrogen-bound ERαLBD to form strong condensates (Boija et al., 2018). In contrast, FUS/TAF15 IDRs is able to self-associate to form phase-separated condensates (Kato and McKnight, 2021). Indeed, chimeric proteins with FUS/TAF15 IDR and AR∆IDR formed condensates in some cells even without DHT treatment (Figures 5A and 5B). Therefore, despite the common feature of low complexity for all IDRs, different IDRs have distinct multivalent interaction capacities and properties for their functional specificity. These results further support our hypothesis that the behavior of a hormone receptor IDR is specifically regulated by its own LBD and hormone signaling.

Multiple lines of evidence indicate that polyQ tract expansion within AR can alter its multivalent interaction and phase separation behaviors (Figure 6). First, the saturation concentration for NTDpQ69 was lower than that of NTDwt in the *in vitro* droplet formation assay. Second, in LNCaP cells, DHT-induced GFP-ARpQ69 condensates were less sensitive to 1,6-HD treatment compared to GFP-ARwt. Third, GFP-ARpQ69 condensates in cells showed slower and less complete recovery than GFP-ARwt condensates in the FRAP assay. All these data suggest that polyQ expansion strengthens AR-AR multivalent interactions and are consistent with a recent report that alanine repeats in HOXD13 protein can elevate phase separation (Basu et al., 2020). Previous studies reported that the Leu-rich region preceding the polyQ repeats can turn the polyQ tract into an α-helical structure, preventing AR from aggregating (Eftekharzadeh et al., 2016; Escobedo et al., 2019). Our data suggest that ARpQ69 condensates might be more “gel/solid-like” (Figure 6), and this altered material state resulted in reduced interactions between AR and other enhancer components, such as MED1 (Figures 7D and 7E).

In summary, we show here AR condensation or its multivalent interaction ability can be weakened or strengthened through genetic and chemical approaches. As either disruption (by 1,6-HD, ∆IDR, 7FS mutation, or IDR swapping) or elevation (by polyQ expansion) of AR condensation led to impaired AR transcriptional activity, we propose that an optimal level of AR multivalent interactions is required for its function (see working model in Figure 7G). Our work using AR as an example provides evidence for the importance of maintaining precise levels of multivalent interactions of signal-dependent IDRs to achieve precise hormone-induced enhancer assembly events. Collectively, our results suggest that disruption of the fine-tuned AR protein multivalent interactions might underlie AR-related human pathologies. Therefore, targeting AR condensates might be a potential therapeutic strategy for treating prostate cancer and other AR-involved diseases.

### Limitations of the study

It’s known that the biophysical properties of protein assemblies driven by multivalent interactions can be substantially distinct, varying from highly fluid and liquid-like to more viscous or gel/solid-like (Alberti et al., 2019; Weber, 2017). We show that either reduced (1,6-HD treatment and 7FS mutation) or enhanced multivalent interactions lead to reduced AR transcriptional activity. We further demonstrate that either disruption (∆IDR) or elevation (pQ69) results in reduced interactions between AR and other enhancer components. Therefore, our data suggest that an optimal level of AR multivalent interactions is required for its normal activity. But our study does not address how a specific biophysical status of AR condensates affects its cellular function.

In KD/SBMA, polyQ-AR forms inclusions in various neural and non-neural tissues but it’s unclear whether the inclusions are causal, protective, or unrelated to pathology. It has been proposed that the inclusions might not only cause gain-of-function toxicity but also sequester AR proteins, leading to a loss of normal AR function. Indeed, transcriptional dysregulation is found in cellular and mouse models of KD/SBMA (Monks et al., 2007), but how polyQ expansion regulates AR function in transcription is unclear. Our data suggest that polyQ expansion leads to more “gel/solid-like” AR condensates, resulting in reduced interactions between AR and other enhancer components. Our study thus revealed a potential mechanism by which polyQ expansion reduces AR transcriptional activity in KD/SBMA. However, our study was done in prostate cancer cell lines. It remains to be tested if this enhancer mechanism can be applied to neural and non-neural cells *in vivo*.

## Figure legends

**Figure S1.**
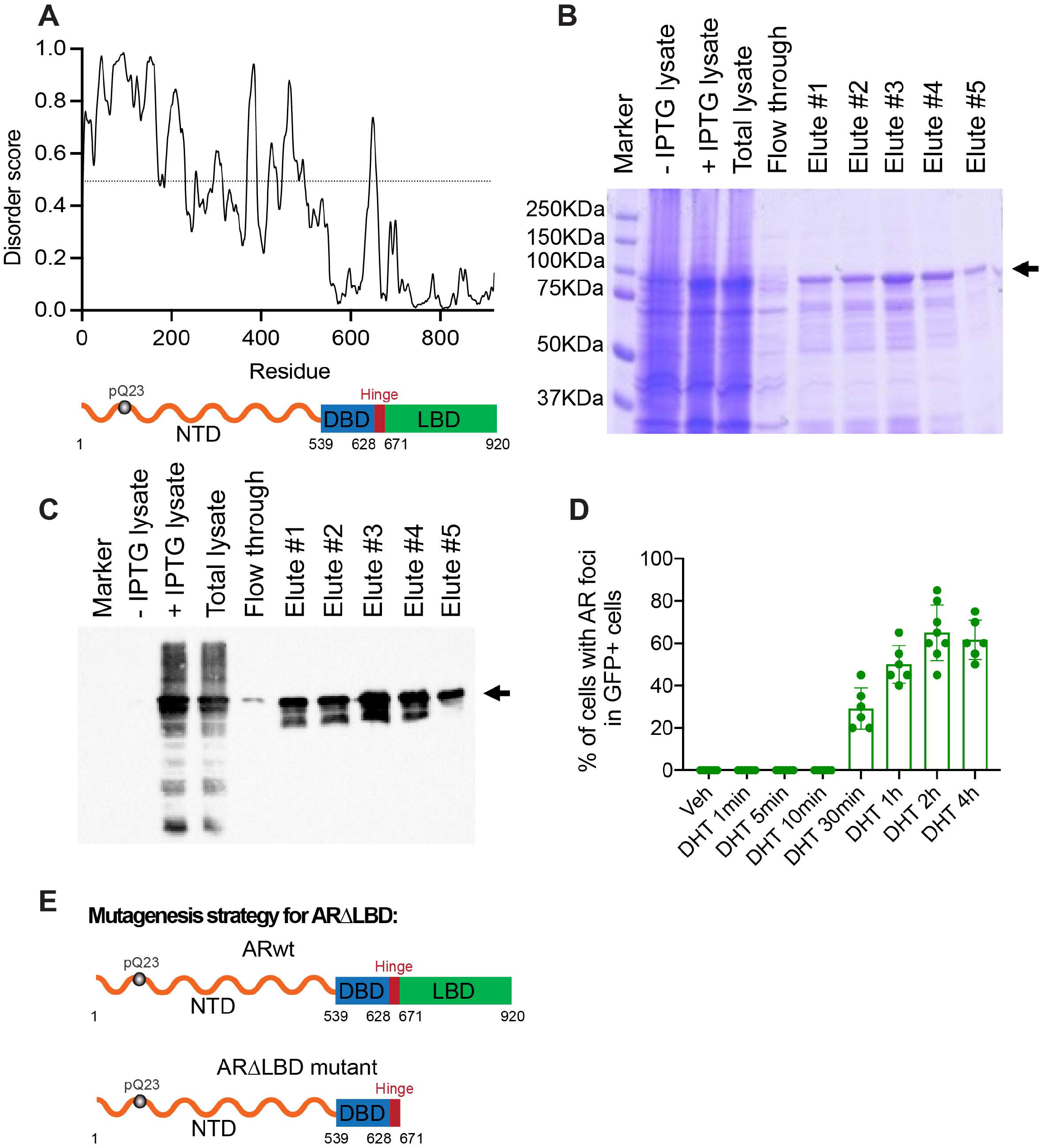
AR NTD is a low complexity domain. **(A)** Plot of intrinsic protein disorder probability for human AR protein using PrDos (http://prdos.hgc.jp/cgi-bin/top.cgi) showing that the NTD has a high disorder score. Different domains of AR protein are indicated at the bottom. **(B)** Coomassie staining of indicated protein samples. Prior to protein purification, we compared cell lysates from cells without and with IPTG induction. We observed an induced band between 75 KDa and 100 KDa, which matched the size of His-GFP-ARNTD (87 KDa). Total lysate from the protein purification experiment, flow through, and elutes from Ni+ columns were also examined. **(C)** Western blot using an antibody recognizing AR NTD (sc-7305, Santa Cruz) on indicated protein samples. **(D)** Quantification of LNCaP cells transfected with GFP-ARwt and treated with DHT (100 nM) at different times. Percentages of cells showing AR foci in GFP-positive cells were plotted. **(E)** Schematic illustration of ARwt and AR∆LBD proteins that were expressed in LNCaP cells for rescue experiments in Figure 1H and 1I.

**Figure S2.**
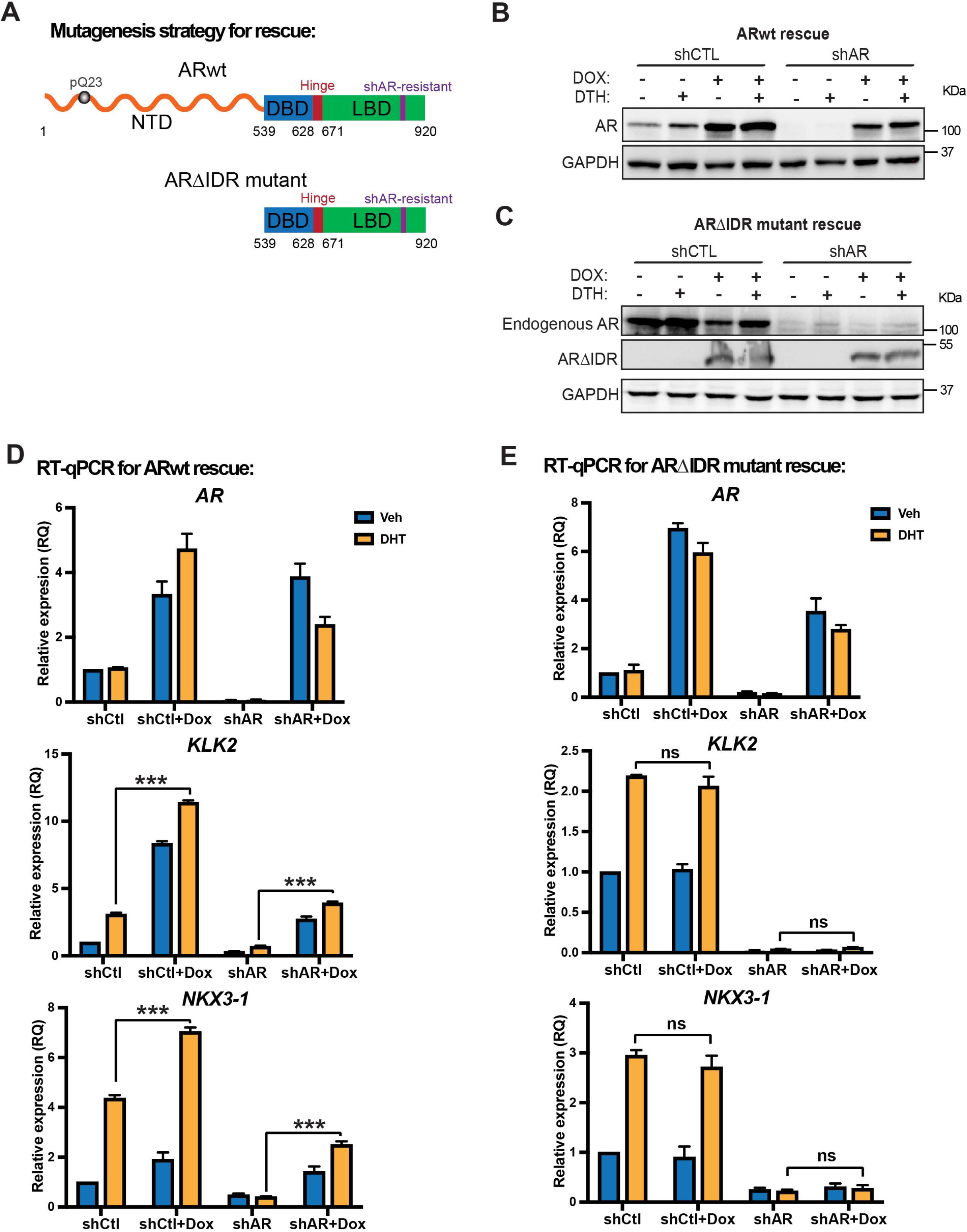
AR IDR is required for AR transcriptional activity. **(A)** Schematic illustration of ARwt and AR∆IDR proteins that were expressed in LNCaP cells for rescue experiments in Figures 2 and S2. **(B)** Western blots to show AR expression levels for the indicated conditions in the rescue experiments with GAPDH as a loading control. LNCaP cells stably expressing Dox-inducible exogenous Arwt were transfected with control or AR shRNA to knock down endogenous AR. shAR effectively knocked down endogenous AR expression and doxycycline-induced exogenous ARwt expression at a level comparable to endogenous AR. **(C)** Western blots to show AR expression levels for the indicated conditions in the rescue experiments with GAPDH as a loading control. LNCaP cells stably expressing Dox-inducible exogenous AR∆IDR were transfected with control or AR shRNA to knock down endogenous AR. Endogenous AR was detected using an antibody recognizing AR N-terminus and exogenous AR∆IDR was detected using an antibody recognizing AR C-terminus. **(D-E)** RT-qPCR data to show the mRNA expression levels of AR and AR target genes (KLK2 and NKX3-1) under the indicated conditions. LNCaP cells stably expressing Dox-inducible exogenous ARwt or AR∆IDR were transfected with control or AR shRNA to knock down endogenous AR. shAR effectively knocked down endogenous AR expression and doxycycline-induced exogenous ARwt or AR∆IDR expression was at a comparable level as endogenous AR. The exogenous ARwt expression was able to substitute for endogenous AR to promote target gene expression. However, exogenous AR∆IDR expression was not able to replace the endogenous AR to promote target gene expression. Statistics: one-way ANOVA, ns: non-significant, ***P < 0.001.

**Figure S3.**
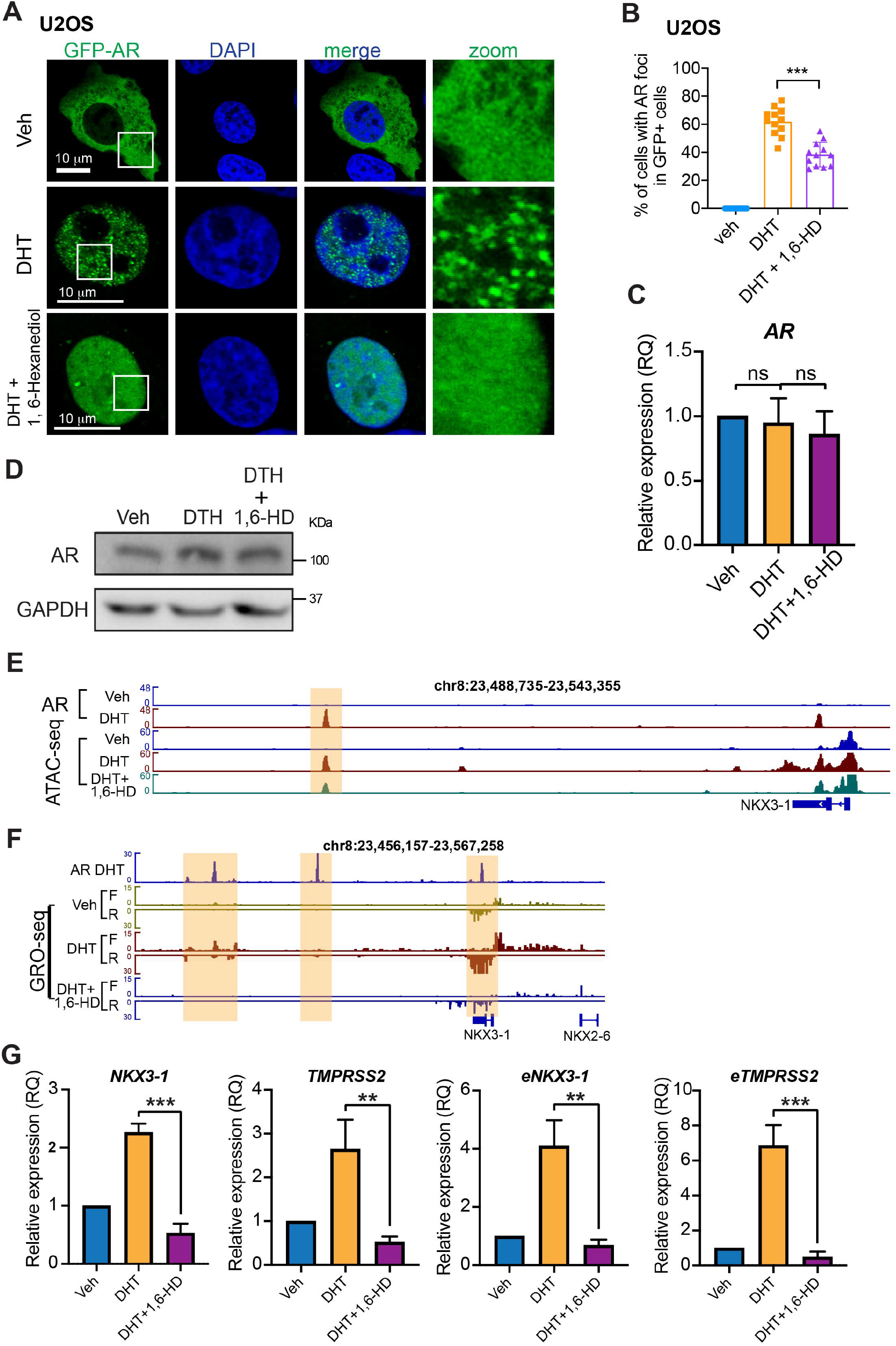
Suppression of AR condensate formation with 1,6-Hexanediol inhibits AR transcriptional activity. **(A-B)** Representative confocal images and quantification of U2OS cells transiently transfected with GFP-ARwt and treated with vehicle, DHT, or DHT+1,6-HD. Like in LNCaP cells, 1,6-HD treatment significantly disrupted DHT-induced AR foci formation. White boxes indicate the zoomed regions shown on the right. Scale bar: 10 µm. Percentages of cells showing AR foci in GFP-positive cells were plotted. Statistics: one-way ANOVA, ***P < 0.001. **(C, G)** RT-qPCR data to show the mRNA expression levels of AR and AR target genes (KLK2 and NKX3-1), as well as the enhancer RNA expression level of AR target genes, under the indicated conditions. 1,6-HD treatment did not affect AR mRNA expression. However, it significantly reduced both mRNA and eRNA levels of AR target genes. Statistics: one-way ANOVA, ns: non-significant, ***P < 0.001, **P < 0.01. **(D)** Western blots to show AR expression levels for the indicated conditions. 1,6-HD treatment did not affect AR protein level. **(E)** Genome browser view of ATAC-seq signals on the enhancers of AR target gene NKX3-1 (highlighted by light yellow color). 1,6-HD treatment reduced DHT-induced chromatin opening on NKX3-1 enhancer. **(F)** Genome browser view of GRO-seq signals on the enhancers of AR target gene NKX3-1 (highlighted by light yellow color). 1,6-HD treatment reduced DHT-induced mRNA and eRNA transcription.

**Figure S4.**
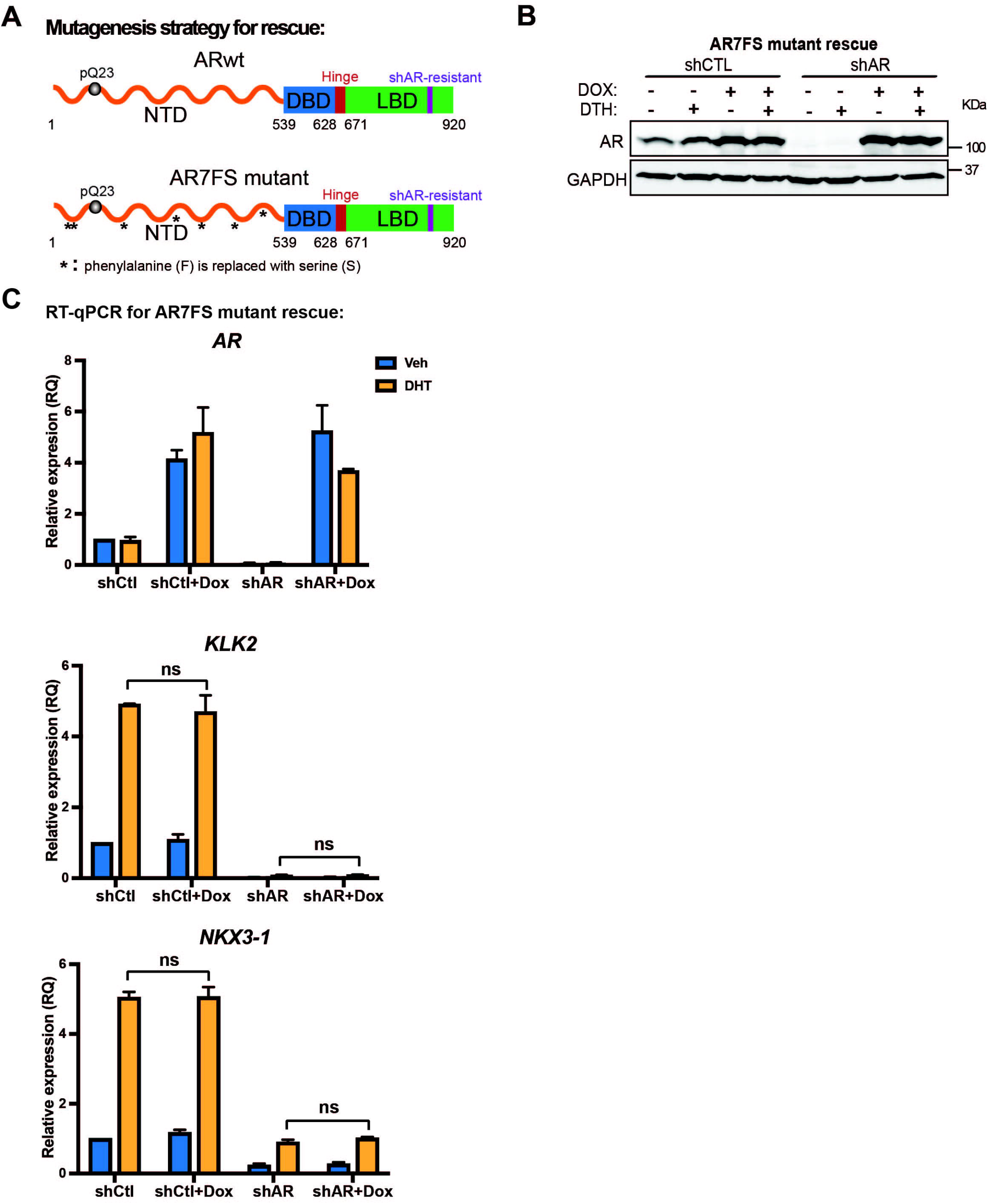
Aromatic residues within AR IDR are required for AR multivalent interactions and transcriptional activity. **(A)** Schematic illustration of ARwt and AR7FS proteins that were expressed in LNCaP cells for rescue experiments in Figures 4 and S4. **(B)** Western blots to show AR expression levels for the indicated conditions in the rescue experiments with GAPDH as a loading control. LNCaP cells stably expressing Dox-inducible exogenous AR7FS were transfected with control or AR shRNA to knock down endogenous AR. shAR effectively knocked down endogenous AR expression and doxycycline-induced exogenous AR7FS expression at a level comparable to endogenous AR. **(C)** RT-qPCR data to show the mRNA expression levels of AR and AR target genes (KLK2 and NKX3-1) under the indicated conditions. shAR effectively knocked down endogenous AR expression and doxycycline-induced exogenous AR7FS expression was at a comparable level as endogenous AR. However, exogenous AR7FS expression was not able to substitute for endogenous AR to promote target gene expression. Statistics: one-way ANOVA, ns: non-significant.

**Figure S5.**
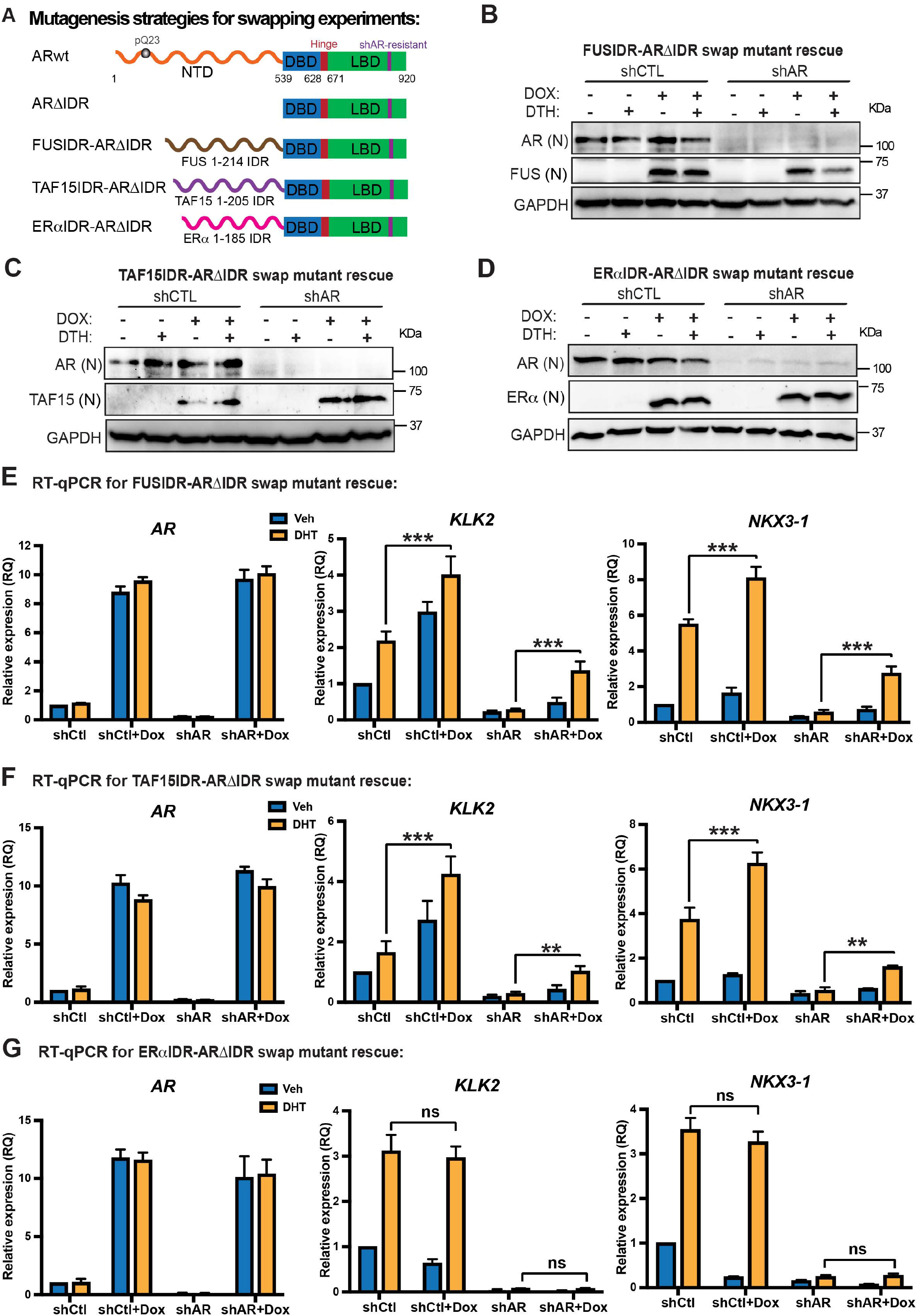
AR IDR can be functionally substituted by selective IDRs. **(A)** Schematic illustration of ARwt and swapped AR mutant proteins that were expressed in LNCaP cells for rescue experiments in Figures 5 and S5. **(B-D)** Western blots to show the expression levels of swapped AR fusion proteins at the indicated conditions with GAPDH as a loading control. LNCaP cells stably expressing Dox-inducible exogenous fusion proteins (FUSIDR-AR∆IDR, TAF15IDR-AR∆IDR, or ERαIDR-AR∆IDR) were transfected with control or AR shRNA to knock down endogenous AR. The antibody recognizing AR N-terminus was used to examine endogenous AR expression, and the antibodies recognizing FUSIDR, TAF15IDR, and ERαIDR were used to examine the expression of fusion proteins. **(E-G)** RT-qPCR data to show the mRNA expression levels of AR target genes (KLK2 and NKX3-1) under the indicated conditions. Expression of FUSIDR-AR∆IDR or TAF15IDR-AR∆IDR, but not ERαIDR-AR∆IDR was sufficient to substitute for endogenous AR to promote DHT-induced AR target gene transcription. Statistics: one-way ANOVA, ns: non-significant, ***P < 0.001, **P < 0.01.

**Figure S6.**
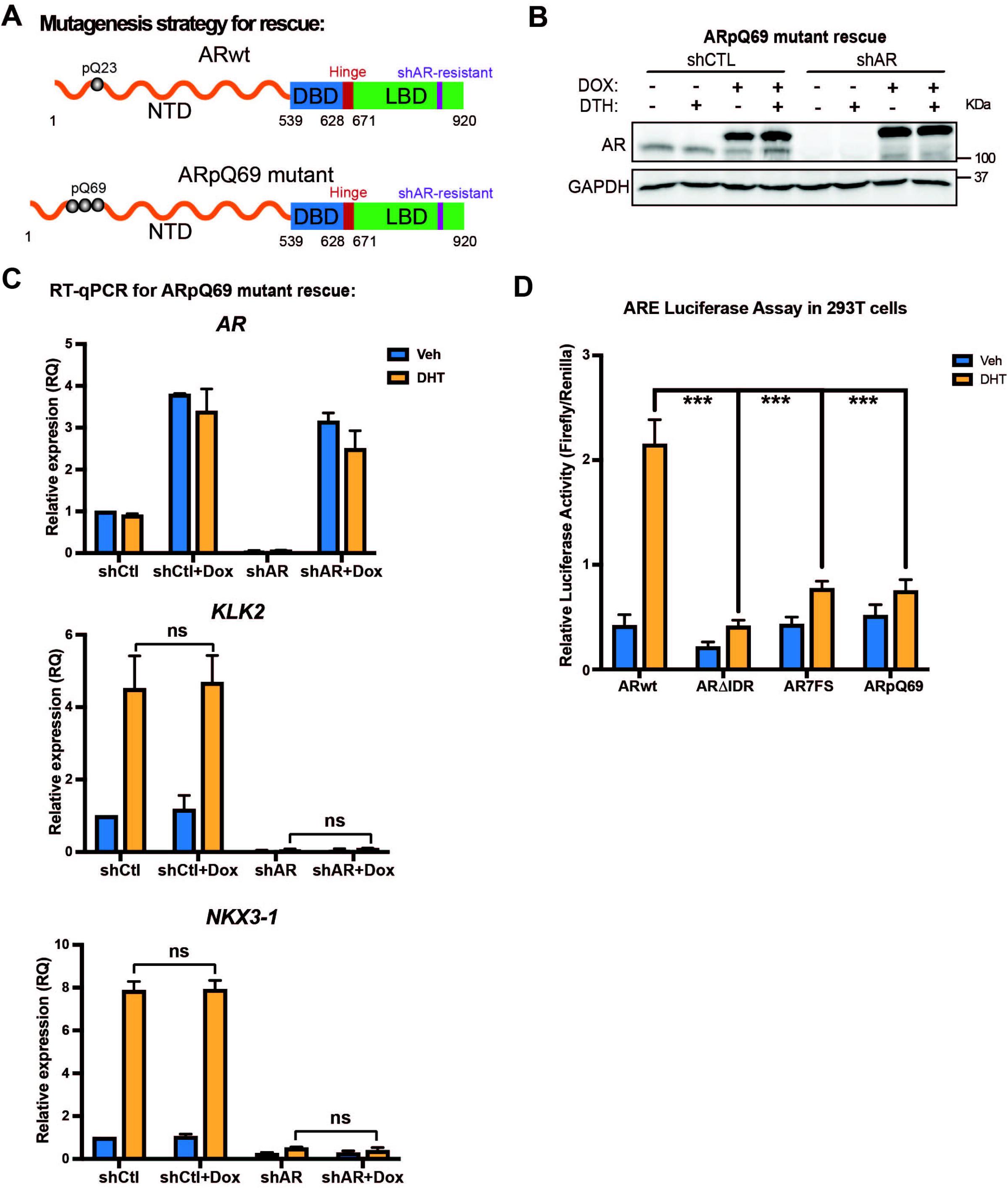
PolyQ tract expansion abolished AR transcriptional activity. **(A)** Schematic illustration of ARwt and ARpQ69 proteins that were expressed in LNCaP cells for rescue experiments in Figures 6 and S6. **(B)** Western blots to show AR expression levels for the indicated conditions in the rescue experiments with GAPDH as a loading control. LNCaP cells stably expressing Dox-inducible exogenous ARpQ69 were transfected with control or AR shRNA to knock down endogenous AR. shAR effectively knocked down endogenous AR expression and doxycycline-induced exogenous ARpQ69 expression at a level comparable to endogenous AR. **(C)** RT-qPCR data to show the mRNA expression levels of AR and AR target genes (KLK2 and NKX3-1) under the indicated conditions. shAR effectively knocked down endogenous AR expression and doxycycline-induced exogenous ARpQ69 expression was at a comparable level as endogenous AR. Exogenous ARpQ69 expression was not able to substitute for endogenous AR to promote target gene expression. Statistics: one-way ANOVA, ns: non-significant. **(D)** Luciferase reporter assay to examine the transcriptional activity of the indicated AR proteins and conditions. 293T cells were co-transfected with the luciferase reporter vector containing 3xARE and one of the AR constructs followed by luciferase activity measurement. Compared to ARwt, AR∆IDR, AR7FS, and ARpQ69 all showed significant reduction in transcriptional activity. Statistics: one-way ANOVA, ns: non-significant, ***P < 0.001.

**Figure S7.**
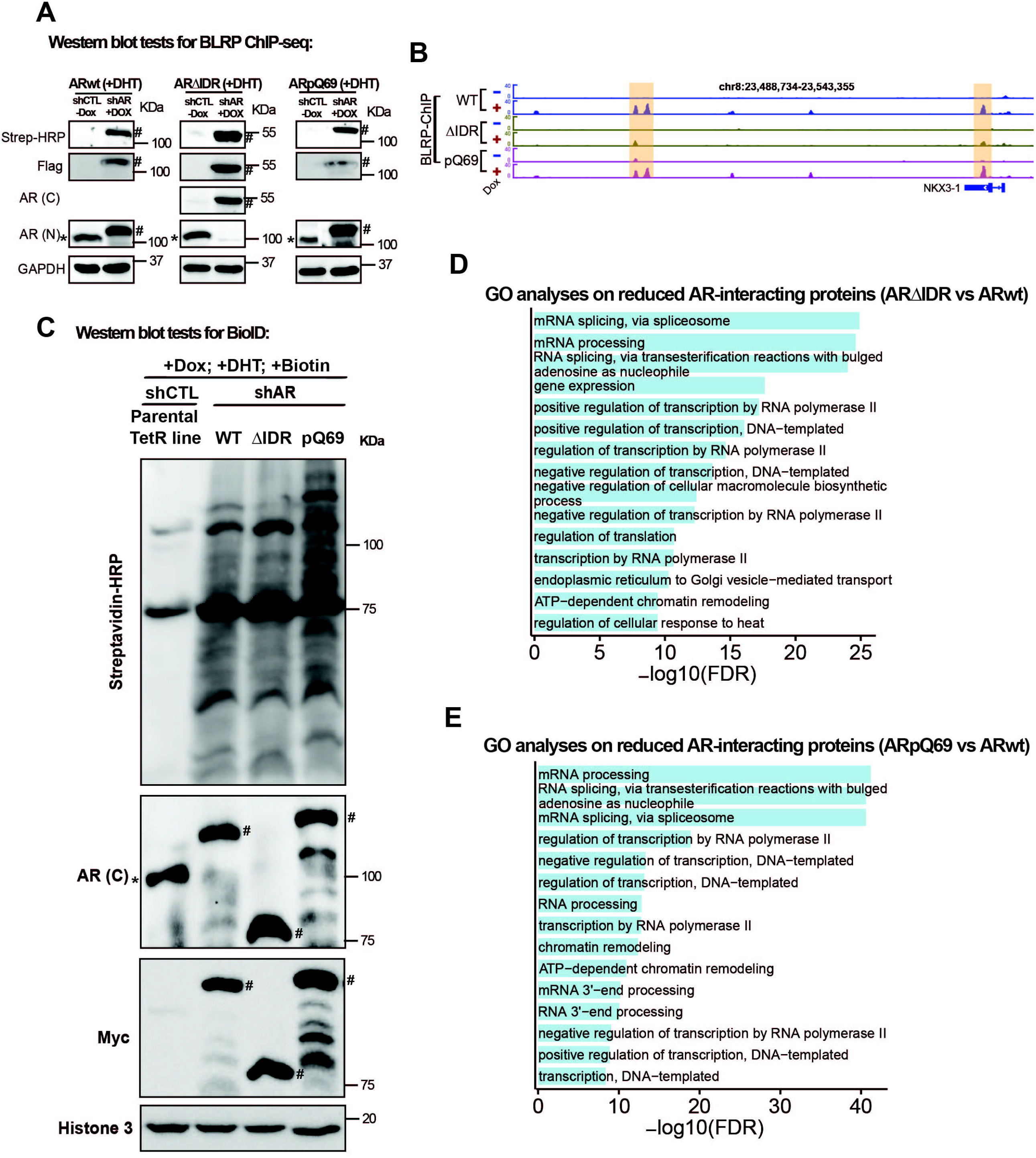
An optimal level of AR multivalent interactions is required for enhancer assembly. **(A)** Western blots to show AR expression levels for the indicated conditions in the ChIP-seq experiment with GAPDH as the loading control. Endogenous AR was knocked down with shRNA and exogenous expression of BLRP-tagged AR protein was induced by doxycycline. Streptavidin-HRP and Flag western blots are to show the expression of exogenous BLRP-tagged AR proteins (wt, ∆IDR and pQ69). AR antibody recognizing C-terminus was used to detect AR∆IDR, and AR antibody recognizing N-terminus was used to detect both endogenous AR and exogenous (wt and pQ69) proteins. The endogenous AR was labeled with * and the BLRP-tagged exogenous AR was labeled with #. **(B)** Genome browser view of ChIP-seq signals on the enhancers of AR target gene NKX3-1 (highlighted by light yellow color) showing that IDR deletion diminished AR binding but polyQ track expansion did not affect AR binding on AR enhancers. **(C)** Western blots confirming the inducible expression and *in vivo* proximity biotinylation in the established tet-on stable LNCaP cell lines for AR BioID (wt, ∆IDR, and pQ69). The fractionation of nuclear fractions and loading amount were confirmed with Western blots for Histone H3 (nucleus-specific marker). Endogenous AR was knocked down with shRNA, which was confirmed with an antibody recognizing AR C-terminus. The doxycycline-induced Myc-BirA*-AR fusion protein expression was detected by antibodies recognizing AR C-terminus and Myc respectively (the endogenous AR was labeled with * and the tagged exogenous AR was labeled with #). Various proteins biotinylated by Myc-BirA*-AR were detected using streptavidin-HRP blot. **(D-E)** Gene Ontology analyses on AR-interacting proteins that showed reduced interactions with AR∆IDR (D) or ARpQ69 (E) compared to ARwt. Proteins involved in mRNA processing and RNA pol II transcription were the top-ranked enriched categories.

## Acknowledgements

Z.L. is a CPRIT Scholar in Cancer Research. This work was supported by funds from CPRIT RR160017 to Z.L., V Foundation V2016-017 to Z.L., V Foundation DVP2019-018 to Z.L., Voelcker Fund Young Investigator Award to Z.L., UT Rising STARs Award to Z.L., Susan G. Komen CCR Award CCR17483391 to Z.L., NCI U54 CA217297/PRJ001 to Z.L., the Mary Kay Foundation Cancer Research Grant to Z.L., Voelcker Fund Young Investigator Award to L.C., NIA R01AG070214 to L.C., and NIA R01AG071591 to L.C., Pew-Stewart Scholar Award to S.C., Searle Scholar Award to S.C., the Shurl and Kay Curci Foundation Research Grant to S.C., and Merkin Innovation Seed Grant for S.C.. Research reported in this publication was also supported by the NIGMS of the NIH under Award Number R01GM137009 to Z. Liu. The content is solely the responsibility of the authors and does not necessarily represent the official views of the NIH. We would like to thank all the members of Liu and Chen labs for technical assistance and helpful discussion, especially for Dr. Mingjun Bi for his technical support on ATAC-seq and ChIP-seq and Dr. Jian Gao for his help with imaging and 1,6 HD treatment work.

## Author Contributions

Z.L., L.C., and S.C. conceived the work and designed the study. L.C., Z.L., Q.H., L.R., and E.Z. performed experiments and data analyses with assistance from R.A., X.L., S.K., P.X., and E.S.. Z.Z. performed the computational analyses for all next-generation sequencing assays. Z.L. supervised the research and oversaw the project. L.C. and Z.L. wrote the manuscript with input from all authors. K.X., Q.W., and T. H. provided comments and reviewed the manuscript.

## References

Alberti, S., Gladfelter, A., and Mittag, T. (2019). Considerations and Challenges in Studying Liquid-Liquid Phase Separation and Biomolecular Condensates. Cell 176, 419–434.

Banani, S.F., Lee, H.O., Hyman, A.A., and Rosen, M.K. (2017). Biomolecular condensates: organizers of cellular biochemistry. Nat Rev Mol Cell Biol 18, 285–298.

Basu, S., Mackowiak, S.D., Niskanen, H., Knezevic, D., Asimi, V., Grosswendt, S., Geertsema, H., Ali, S., Jerkovic, I., Ewers, H., et al. (2020). Unblending of Transcriptional Condensates in Human Repeat Expansion Disease. Cell 181, 1062–1079 e1030.

Bi, M., Zhang, Z., Jiang, Y.Z., Xue, P., Wang, H., Lai, Z., Fu, X., De Angelis, C., Gong, Y., Gao, Z., et al. (2020). Enhancer reprogramming driven by high-order assemblies of transcription factors promotes phenotypic plasticity and breast cancer endocrine resistance. Nat Cell Biol 22, 701–715.

Boehning, M., Dugast-Darzacq, C., Rankovic, M., Hansen, A.S., Yu, T., Marie-Nelly, H., McSwiggen, D.T., Kokic, G., Dailey, G.M., Cramer, P., et al. (2018). RNA polymerase II clustering through carboxy-terminal domain phase separation. Nat Struct Mol Biol 25, 833–840.

Boija, A., Klein, I.A., Sabari, B.R., Dall’Agnese, A., Coffey, E.L., Zamudio, A.V., Li, C.H., Shrinivas, K., Manteiga, J.C., Hannett, N.M., et al. (2018). Transcription Factors Activate Genes through the Phase-Separation Capacity of Their Activation Domains. Cell 175, 1842–1855 e1816.

Brodsky, S., Jana, T., Mittelman, K., Chapal, M., Kumar, D.K., Carmi, M., and Barkai, N. (2020). Intrinsically Disordered Regions Direct Transcription Factor In Vivo Binding Specificity. Mol Cell 79, 459–471 e454.

Bugaj, L.J., Choksi, A.T., Mesuda, C.K., Kane, R.S., and Schaffer, D.V. (2013). Optogenetic protein clustering and signaling activation in mammalian cells. Nat Methods 10, 249–252.

Cho, W.K., Spille, J.H., Hecht, M., Lee, C., Li, C., Grube, V., and Cisse, II (2018). Mediator and RNA polymerase II clusters associate in transcription-dependent condensates. Science 361, 412–415.

Choi, J.M., Holehouse, A.S., and Pappu, R.V. (2020). Physical Principles Underlying the Complex Biology of Intracellular Phase Transitions. Annu Rev Biophys 49, 107–133.

Chong, S., Dugast-Darzacq, C., Liu, Z., Dong, P., Dailey, G.M., Cattoglio, C., Heckert, A., Banala, S., Lavis, L., Darzacq, X., et al. (2018). Imaging dynamic and selective low-complexity domain interactions that control gene transcription. Science 361.

Chong, S., Graham, T.G.W., Dugast-Darzacq, C., Dailey, G.M., Darzacq, X., and Tjian, R. (2022). Tuning levels of low-complexity domain interactions to modulate endogenous oncogenic transcription. Mol Cell 82, 2084–2097 e2085.

Decker, C.J., Teixeira, D., and Parker, R. (2007). Edc3p and a glutamine/asparagine-rich domain of Lsm4p function in processing body assembly in Saccharomyces cerevisiae. J Cell Biol 179, 437–449.

Eftekharzadeh, B., Piai, A., Chiesa, G., Mungianu, D., Garcia, J., Pierattelli, R., Felli, I.C., and Salvatella, X. (2016). Sequence Context Influences the Structure and Aggregation Behavior of a PolyQ Tract. Biophys J 110, 2361–2366.

Escobedo, A., Topal, B., Kunze, M.B.A., Aranda, J., Chiesa, G., Mungianu, D., Bernardo-Seisdedos, G., Eftekharzadeh, B., Gairi, M., Pierattelli, R., et al. (2019). Side chain to main chain hydrogen bonds stabilize a polyglutamine helix in a transcription factor. Nat Commun 10, 2034.

Gilks, N., Kedersha, N., Ayodele, M., Shen, L., Stoecklin, G., Dember, L.M., and Anderson, P. (2004). Stress granule assembly is mediated by prion-like aggregation of TIA-1. Mol Biol Cell 15, 5383–5398.

Gottlieb, B., Beitel, L.K., Nadarajah, A., Paliouras, M., and Trifiro, M. (2012). The androgen receptor gene mutations database: 2012 update. Hum Mutat 33, 887–894.

Hah, N., Murakami, S., Nagari, A., Danko, C.G., and Kraus, W.L. (2013). Enhancer transcripts mark active estrogen receptor binding sites. Genome Res 23, 1210–1223.

Hazelett, D.J., Rhie, S.K., Gaddis, M., Yan, C., Lakeland, D.L., Coetzee, S.G., Ellipse, G.-O.N.c., Practical, c., Henderson, B.E., Noushmehr, H., et al. (2014). Comprehensive functional annotation of 77 prostate cancer risk loci. PLoS Genet 10, e1004102.

He, B., Gampe, R.T., Jr., Kole, A.J., Hnat, A.T., Stanley, T.B., An, G., Stewart, E.L., Kalman, R.I., Minges, J.T., and Wilson, E.M. (2004). Structural basis for androgen receptor interdomain and coactivator interactions suggests a transition in nuclear receptor activation function dominance. Mol Cell 16, 425–438.

Heery, D.M., Hoare, S., Hussain, S., Parker, M.G., and Sheppard, H. (2001). Core LXXLL motif sequences in CREB-binding protein, SRC1, and RIP140 define affinity and selectivity for steroid and retinoid receptors. J Biol Chem 276, 6695–6702.

Hnisz, D., Shrinivas, K., Young, R.A., Chakraborty, A.K., and Sharp, P.A. (2017). A Phase Separation Model for Transcriptional Control. Cell 169, 13–23.

Ho, W.L., and Huang, J.R. (2022). The return of the rings: Evolutionary convergence of aromatic residues in the intrinsically disordered regions of RNA-binding proteins for liquid-liquid phase separation. Protein Sci 31, e4317.

Janicki, S.M., Tsukamoto, T., Salghetti, S.E., Tansey, W.P., Sachidanandam, R., Prasanth, K.V., Ried, T., Shav-Tal, Y., Bertrand, E., Singer, R.H., et al. (2004). From silencing to gene expression: real-time analysis in single cells. Cell 116, 683–698.

Jenster, G., van der Korput, H.A., Trapman, J., and Brinkmann, A.O. (1995). Identification of two transcription activation units in the N-terminal domain of the human androgen receptor. J Biol Chem 270, 7341–7346.

Kato, M., and McKnight, S.L. (2018). A Solid-State Conceptualization of Information Transfer from Gene to Message to Protein. Annu Rev Biochem 87, 351–390.

Kato, M., and McKnight, S.L. (2021). The low-complexity domain of the FUS RNA binding protein self-assembles via the mutually exclusive use of two distinct cross-beta cores. Proc Natl Acad Sci U S A 118.

Kwon, I., Kato, M., Xiang, S., Wu, L., Theodoropoulos, P., Mirzaei, H., Han, T., Xie, S., Corden, J.L., and McKnight, S.L. (2013). Phosphorylation-regulated binding of RNA polymerase II to fibrous polymers of low-complexity domains. Cell 155, 1049–1060.

Lam, M.T., Li, W., Rosenfeld, M.G., and Glass, C.K. (2014). Enhancer RNAs and regulated transcriptional programs. Trends Biochem Sci 39, 170–182.

Li, H.R., Chiang, W.C., Chou, P.C., Wang, W.J., and Huang, J.R. (2018). TAR DNA-binding protein 43 (TDP-43) liquid-liquid phase separation is mediated by just a few aromatic residues. J Biol Chem 293, 6090–6098.

Lin, Y., Currie, S.L., and Rosen, M.K. (2017). Intrinsically disordered sequences enable modulation of protein phase separation through distributed tyrosine motifs. J Biol Chem 292, 19110–19120.

Lin, Y., Mori, E., Kato, M., Xiang, S., Wu, L., Kwon, I., and McKnight, S.L. (2016). Toxic PR Poly-Dipeptides Encoded by the C9orf72 Repeat Expansion Target LC Domain Polymers. Cell 167, 789–802 e712.

Liu, Z., Merkurjev, D., Yang, F., Li, W., Oh, S., Friedman, M.J., Song, X., Zhang, F., Ma, Q., Ohgi, K.A., et al. (2014). Enhancer activation requires trans-recruitment of a mega transcription factor complex. Cell 159, 358–373.

Lu, H., Yu, D., Hansen, A.S., Ganguly, S., Liu, R., Heckert, A., Darzacq, X., and Zhou, Q. (2018). Phase-separation mechanism for C-terminal hyperphosphorylation of RNA polymerase II. Nature 558, 318–323.

Malinen, M., Niskanen, E.A., Kaikkonen, M.U., and Palvimo, J.J. (2017). Crosstalk between androgen and pro-inflammatory signaling remodels androgen receptor and NF-kappaB cistrome to reprogram the prostate cancer cell transcriptome. Nucleic Acids Res 45, 619–630.

Matias, P.M., Donner, P., Coelho, R., Thomaz, M., Peixoto, C., Macedo, S., Otto, N., Joschko, S., Scholz, P., Wegg, A., et al. (2000). Structural evidence for ligand specificity in the binding domain of the human androgen receptor. Implications for pathogenic gene mutations. J Biol Chem 275, 26164–26171.

McCampbell, A., Taye, A.A., Whitty, L., Penney, E., Steffan, J.S., and Fischbeck, K.H. (2001). Histone deacetylase inhibitors reduce polyglutamine toxicity. Proc Natl Acad Sci U S A 98, 15179–15184.

McCampbell, A., Taylor, J.P., Taye, A.A., Robitschek, J., Li, M., Walcott, J., Merry, D., Chai, Y., Paulson, H., Sobue, G., et al. (2000). CREB-binding protein sequestration by expanded polyglutamine. Hum Mol Genet 9, 2197–2202.

McSwiggen, D.T., Mir, M., Darzacq, X., and Tjian, R. (2019). Evaluating phase separation in live cells: diagnosis, caveats, and functional consequences. Genes Dev 33, 1619–1634.

Mittag, T., and Pappu, R.V. (2022). A conceptual framework for understanding phase separation and addressing open questions and challenges. Mol Cell 82, 2201–2214.

Monaghan, A.E., and McEwan, I.J. (2016). A sting in the tail: the N-terminal domain of the androgen receptor as a drug target. Asian J Androl 18, 687–694.

Monks, D.A., Johansen, J.A., Mo, K., Rao, P., Eagleson, B., Yu, Z., Lieberman, A.P., Breedlove, S.M., and Jordan, C.L. (2007). Overexpression of wild-type androgen receptor in muscle recapitulates polyglutamine disease. Proc Natl Acad Sci U S A 104, 18259–18264.

Nair, S.J., Yang, L., Meluzzi, D., Oh, S., Yang, F., Friedman, M.J., Wang, S., Suter, T., Alshareedah, I., Gamliel, A., et al. (2019). Phase separation of ligand-activated enhancers licenses cooperative chromosomal enhancer assembly. Nat Struct Mol Biol 26, 193–203.

Panigrahi, A., and O’Malley, B.W. (2021). Mechanisms of enhancer action: the known and the unknown. Genome Biol 22, 108.

Peskett, T.R., Rau, F., O’Driscoll, J., Patani, R., Lowe, A.R., and Saibil, H.R. (2018). A Liquid to Solid Phase Transition Underlying Pathological Huntingtin Exon1 Aggregation. Mol Cell 70, 588–601 e586.

Plank, J.L., and Dean, A. (2014). Enhancer function: mechanistic and genome-wide insights come together. Molecular cell 55, 5–14.

Portz, B., Lee, B.L., and Shorter, J. (2021). FUS and TDP-43 Phases in Health and Disease. Trends Biochem Sci 46, 550–563.

Roux, K.J., Kim, D.I., Raida, M., and Burke, B. (2012). A promiscuous biotin ligase fusion protein identifies proximal and interacting proteins in mammalian cells. The Journal of cell biology 196, 801–810.

Russo, J.W., Nouri, M., and Balk, S.P. (2019). Androgen Receptor Interaction with Mediator Complex Is Enhanced in Castration-Resistant Prostate Cancer by CDK7 Phosphorylation of MED1. Cancer Discov 9, 1490–1492.

Sabari, B.R., Dall’Agnese, A., Boija, A., Klein, I.A., Coffey, E.L., Shrinivas, K., Abraham, B.J., Hannett, N.M., Zamudio, A.V., Manteiga, J.C., et al. (2018). Coactivator condensation at super-enhancers links phase separation and gene control. Science 361.

Saporita, A.J., Zhang, Q., Navai, N., Dincer, Z., Hahn, J., Cai, X., and Wang, Z. (2003). Identification and characterization of a ligand-regulated nuclear export signal in androgen receptor. J Biol Chem 278, 41998–42005.

Shin, Y., Berry, J., Pannucci, N., Haataja, M.P., Toettcher, J.E., and Brangwynne, C.P. (2017). Spatiotemporal Control of Intracellular Phase Transitions Using Light-Activated optoDroplets. Cell 168, 159–171 e114.

Simental, J.A., Sar, M., Lane, M.V., French, F.S., and Wilson, E.M. (1991). Transcriptional activation and nuclear targeting signals of the human androgen receptor. J Biol Chem 266, 510–518.

Stelloo, S., Bergman, A.M., and Zwart, W. (2019). Androgen receptor enhancer usage and the chromatin regulatory landscape in human prostate cancers. Endocr Relat Cancer 26, R267–R285.

Stelloo, S., Nevedomskaya, E., Kim, Y., Hoekman, L., Bleijerveld, O.B., Mirza, T., Wessels, L.F.A., van Weerden, W.M., Altelaar, A.F.M., Bergman, A.M., et al. (2018). Endogenous androgen receptor proteomic profiling reveals genomic subcomplex involved in prostate tumorigenesis. Oncogene 37, 313–322.

Tan, M.H., Li, J., Xu, H.E., Melcher, K., and Yong, E.L. (2015). Androgen receptor: structure, role in prostate cancer and drug discovery. Acta Pharmacol Sin 36, 3–23.

Thomas, M., Yu, Z., Dadgar, N., Varambally, S., Yu, J., Chinnaiyan, A.M., and Lieberman, A.P. (2005). The unfolded protein response modulates toxicity of the expanded glutamine androgen receptor. J Biol Chem 280, 21264–21271.

Trojanowski, J., Frank, L., Rademacher, A., Mucke, N., Grigaitis, P., and Rippe, K. (2022). Transcription activation is enhanced by multivalent interactions independent of phase separation. Mol Cell 82, 1878–1893 e1810.

van Royen, M.E., Cunha, S.M., Brink, M.C., Mattern, K.A., Nigg, A.L., Dubbink, H.J., Verschure, P.J., Trapman, J., and Houtsmuller, A.B. (2007). Compartmentalization of androgen receptor protein-protein interactions in living cells. J Cell Biol 177, 63–72.

Wan, L., Chong, S., Xuan, F., Liang, A., Cui, X., Gates, L., Carroll, T.S., Li, Y., Feng, L., Chen, G., et al. (2020). Impaired cell fate through gain-of-function mutations in a chromatin reader. Nature 577, 121–126.

Wang, J., Choi, J.M., Holehouse, A.S., Lee, H.O., Zhang, X., Jahnel, M., Maharana, S., Lemaitre, R., Pozniakovsky, A., Drechsel, D., et al. (2018). A Molecular Grammar Governing the Driving Forces for Phase Separation of Prion-like RNA Binding Proteins. Cell 174, 688–699 e616.

Weber, S.C. (2017). Sequence-encoded material properties dictate the structure and function of nuclear bodies. Curr Opin Cell Biol 46, 62–71.

Wei, M.T., Chang, Y.C., Shimobayashi, S.F., Shin, Y., Strom, A.R., and Brangwynne, C.P. (2020). Nucleated transcriptional condensates amplify gene expression. Nat Cell Biol 22, 1187–1196.

Zhu, C., Li, L., Zhang, Z., Bi, M., Wang, H., Su, W., Hernandez, K., Liu, P., Chen, J., Chen, M., et al. (2019). A Non-canonical Role of YAP/TEAD Is Required for Activation of Estrogen-Regulated Enhancers in Breast Cancer. Mol Cell 75, 791–806 e798.

