## Supplemental Method for "Hormone-induced enhancer assembly requires an optimal level of hormone receptor multivalent interactions"

### Materials and Methods

#### Cell culture

LNCaP, U2OS, and 293T cell lines were obtained from ATCC. LNCaP cells were cultured in Roswell Park Memorial Institute (RPMI) 1640 Medium supplemented with 10% FBS and penicillin/streptomycin. U2OS and 293T cells were cultured in Dulbecco's Modified Eagle's Medium (DMEM) supplemented with 10% FBS and penicillin/streptomycin. All cell lines were grown in a humidified incubator with 5% $CO_2$. For DHT stimulation, cells were hormone stripped for 3 days in phenol-free media with 5% charcoal-stripped FBS before receiving 100nM DHT (Sigma) or ethanol vehicle control treatment for 2 hours for androgen signaling induction. We used 1,6 hexanediol (Sigma) to chemically disrupt phase separation in cells. 1,6 hexanediol (1,6 HD) was used at 2.5% in the medium for LNCaP or U2OS cells for different treatment timing for different experiments (5 minutes for immunofluorescence imaging and 2 hours for western blot/RT-qPCR/ATAC-seq/GRO-seq). To study the interactions of AR-AR and AR-MED1 interactions on the chromatin, U2OS 2-6-3 line, which has 200 copies of p3216PECMS2beta plasmid stable insertion at one genome locus (Chong et al., 2018; Janicki et al., 2004). Each copy of p3216PECMS2beta plasmid contains 256 copies of Lac O. Thus, the total Lac O copies in U2OS 2-6-3 line should be 200 x 256 = ~50,000. The culture condition of the U2OS 2-6-3 line is the same as the regular U2OS line that was described above in addition to using 100ug/ml hygromycin for stable selection. All cell lines were routinely tested to ensure they are free of mycoplasma contamination (VenorTMGeM Mycoplasma Detection Kit, Sigma-Aldrich).

#### Protein purification and *in vitro* droplet formation assay

Protein purification was performed as previously described (Sabari et al., 2018) with some modifications. Constructs used for protein purification were generated through gateway LR recombination between entry clones containing AR cDNA and destination vectors modified from pET-mCherry-MED1 or pET-GFP-MED1 (gift from Richard Young Lab). *E. coli* Rosetta (DE3) competent cells were used as the host strain for protein expression. The cells were transformed with various AR plasmids and protein expression was induced with 0.1 mM IPTG in Luria broth (LB) containing 1% glucose, 1X Kanamycin, and 0.5X chloramphenicol. The culture was grown overnight at 18$^{o}$C and pelleted at 3000 rpm for 10 minutes and either proceeded fresh to lysis or stored at -80$^{o}$C. The cells were lysed in a Buffer A (50 mM Tris-HCl, pH 7.5, 500 mM NaCl, 10 mM imidazole, protease inhibitor cocktail (Roche)) and subjected to sonication. The total cell lysate was centrifuged at 20000 rpm for 30 minutes at 4 °C, and the soluble fraction was loaded onto a Ni–NTA agarose resin column that was pre-equilibrated with Buffer A. The resin-bound protein was washed successively with Buffer A and finally eluted with the elution buffer containing a gradient increase in the concentration of Imidazole (50 mM Tris-HCL, pH 7.5, 500 mM NaCl and 50 mM/100 mM/250 mM Imidazole). The eluted proteins were confirmed by running on a 12% SDS Gel, and the proteins were pooled together and subjected to overnight dialysis (50 mM Tris pH 7.5, 125 mM NaCl, 10% glycerol and 1 mM DTT) at 4$^{o}$C. The dialyzed proteins were concentrated using the 3K centricon (Millipore, Sigma) and either proceeded fresh for the LLPS experiment or aliquoted and flash frozen in liquid nitrogen and stored at -80$^{o}$C.

The purified protein was diluted to 40 μM, 20 μM, 10 μM and 5 μM using the dialysis buffer (50 mM Tris pH 7.5, 125 mM NaCl, 10% glycerol and 1 mM DTT). The protein was mixed in a 1:1 ratio with droplet formation buffer (50 mM Tris pH 7.5, 125 mM NaCl, 10% glycerol, 1 mM DTT, with 20% or 40% PEG8000) and 5 μl of the suspension was loaded onto a hand-made cassette with a clean glass slide, spacers, and a coverslip. The droplets were immediately observed under an upright confocal microscope (Zeiss LSM780) using a 100x/1.4 oil objective. ZEN black edition version 2.3 was used for acquisition. Droplet size quantification was done using the "analyze particles" tool of FIJI with the threshold set as the medium area of all particles detected in each image.

**Fluorescence imaging, OptoDroplets assay, and Fluorescence recovery after photobleaching (FRAP)**

To image wildtype and mutant AR proteins in androgen-responsive cells, LNCaP cells or U2OS cells were transfected with various GFP-AR plasmids. The backbone plasmid was modified from pEGFP-C3 (Clonetech) by including a gateway cassette to generate a destination vector that is compatible with the gateway cloning system. Cells cultured on coverslips and expressing GFP-tagged AR proteins (stripping for 3 days before vehicle or DHT treatment) were fixed using 4% PFA and mounted onto slides using antifade mountant with DAPI (Life Technologies, Carlsbad, CA). Images were acquired at Zeiss LSM780 confocal microscope with a 100x/1.4 oil objective. For quantification, we typically counted the number of cells showing GFP-AR foci in every 20 (or 15) GFP-positive cells and plotted the percentage. Puncta fringe visibility was measured as amplitude/average, i.e. (maximal intensity – minimal intensity)/mean intensity. The intensity was measured using ImageJ by drawing a line crossing the center of a puncta followed by line measurement. The line length was approximately twice of the diameter of the puncta. 5 representative puncta were selected from each cell and a total of 30 puncta were measured.

Immunofluorescence for endogenous AR expression was done as below. Cells grown on coated glass coverslips were fixed in 4% paraformaldehyde in 1xPBS for 10 minutes at room temperature (RT) followed by 1xPBS wash for 5 minutes at RT for 3 times. Cells were permeabilized with 0.5% Triton X-100 in PBS for 20 minutes at RT followed by 3 washes. Cells were then blocked with 5% BSA in PBS containing 0.05% Triton X-100 for 30 minutes at RT followed by incubation of primary antibody (anti-AR, Santa Cruz sc-7305, 1:500 dilution in blocking buffer) at 4°C overnight. After 3 washes, cells were incubated with a secondary antibody (anti-mouse IgG-488, 1:1000 dilution in blocking buffer) for 30 minutes at RT followed by 3 washes. Cells on the coverslips were then mounted onto slides using Antifade Mountant with DAPI (Life Technologies, Carlsbad, CA). Images were acquired at Zeiss LSM780 confocal microscope with a 100x/1.4 oil objective. For quantification, the total area of AR puncta per nucleus was measured using the "analyze particles" tool of FIJI.

OptoDroplet assay was performed as previously described (Shin et al., 2017) with some modifications. OptoDroplet constructs used in this study were derived from the pCRY2PHR-mCherryN1 plasmid (Addgene #26866). The plasmid was modified by including a gateway cassette to generate pCRY2PHR-mCherry-gw that is compatible with the gateway cloning system. 48 hours after the transfection of CRY2 fusion constructs into 293T cells that were cultured on coated glass coverslips, cells were transferred to a culture chamber for live cell imaging using a 63x/1.4 oil objective on Zeiss LSM780 microscope. Cells were imaged by using 561 nm of laser for mCherry signal before and after CRY2 activation, which was done using 2% power of 488 nm laser for 30 seconds.

Fluorescence recovery after photobleaching (FRAP) was done using Zeiss LSM780 confocal microscope. For CRY2 puncta, blue light-induced puncta were bleached using 100% power of 560 nm laser, and their fluorescence recovery was monitored. For DHT-induced GFP-AR puncta,

LNCaP cells were cultured on coated glass coverslips and transferred to a culture chamber for live imaging. FRAP was done using 100% of 488 nm laser power. Fluorescence intensity was measured using FIJI. Background intensity was subtracted, and the relative intensity was normalized to pre-bleaching intensity.

**AR self-association assays with LacO array in U2OS cell line**

Human U2OS 2-6-3 cells containing a LacO array with ~50,000 LacO elements in the genome were used to study AR-AR and AR-MED1 interactions on chromatin. For imaging sample preparation, cells were plated on 70% ethanol-pretreated 18 mm circular No. 1 micro cover glasses (Electron Microscopy Sciences, 72229-01) on 12-Well TC Treated Plates (Genesee, 25-106MP), and were transfected with the target constructs by using Invitrogen™ Lipofectamine 3000 Transfection Reagent (Thermo Scientific L3000015) for 24 hours, followed by fixation with 4% paraformaldehyde (Sigma-Aldrich P6148-500G) for 15 minutes.

Fluorescence images were acquired on a Zeiss LSM 980 laser scanning confocal microscope operated by the Biological Imaging Facility of the Beckman Institute at Caltech, using a 63X, NA 1.4 Plan-Apochromat objective. The pinhole size of the mCherry channel was set to 2.00 AU and EYFP channel was set to 1.00 AU, as the resolution was set to 512 x 512 with a digital zoom of 3.0X and line averaging of 2. Excitation laser sources and emission ranges were 514 nm/508-561 nm (EYFP $\lambda$max=513 nm, $\lambda$em=530 nm, shown as green), and 594 nm/561-693 nm (mCherry $\lambda$max=587 nm, $\lambda$em=610 nm, shown as magenta). Each image was collected as a 3D stack of 25-40 images with a spacing of 0.24 µm in the z-direction between slices. Before acquiring the 2-color fluorescence images, we carefully tuned the emission filters to make sure no bleed-through existed between the two channels and adjusted laser intensity and microscope detector gains to ensure that no pixel in the images was saturated.

We quantified protein-protein interactions from the 2-color images using the following steps. First, we selected the slice (#N) in the EYFP channel z stack where a LacO-associated punctum has the highest fluorescence intensity, located the central pixel of the LacO array, and obtained the radial profiles of fluorescence intensity centering the pixel in both EYFP and mCherry channels. Second, we extracted the EYFP intensity radial profile and estimated the radius of the punctum as the distance from the central pixel at which the derivative of fluorescence intensity to distance first drops to zero. Next, we measured the maximum and peripheral mCherry intensities of the punctum. To average out the intensity noise at the single-pixel level, convolution was applied to image slice #N of the mCherry channel using a 5x5 convolution kernel $J_5$ (all-ones matrix). The intensities of four convoluted pixels surrounding the peak intensity pixel were averaged as $I_{peak}$. Two values on the mCherry intensity radial profile at locations immediately outside the punctum periphery were averaged as $I_{periphery}$. Finally, we calculated the intensity ratio $I_{peak}/I_{periphery}$ as a measure of the mCherry enrichment at the LacO array. A ratio above 1 suggests protein-protein interactions.

**shRNA Lentivirus Package and infection**

Mission shRNA lentiviral plasmid targeting AR (TRCN0000003718) and control shRNA (SHC002 or SHC202) were purchased from Sigma. Knockdown experiments with shRNA lentiviruses were conducted according to the standard lentivirus package and transduction protocols from Addgene. These pLKO-based lentiviral shRNA plasmids were co-transfected with packaging plasmids (psPAX2 and pMD2.G from Addgene) into 293T cells. Lentiviruses were harvested and used for LNCaP cell infection in the rescue experiments below.

**AR rescue experiments in LNCaP cells**

To set up the doxycycline-inducible AR-overexpressing LNCaP cell lines, AR cDNAs for wildtype and different mutants synthesized by GenScript (see the details for these cDNAs in the following section) were individually cloned into pCR8/GW/TOPO and then transferred to the pInducer20 destination vector (pInducer20 was a gift from Stephen Elledge; Addgene #44012) using the Gateway system (Life Technologies), followed by co-transfection with packaging plasmids (psPAX2 and pMD2.G from Addgene) into 293T cells to produce lentiviruses. After lentivirus infection, LNCaP cells were selected by 600 µg/ml G418 (Invitrogen) to set up doxycycline-inducible stable cell lines. For the rescue purpose, all AR cDNAs synthesized by GenScript are resistant to AR shRNA TRCN0000003718 because the shRNA target sequence "CACCAATGTCAACTCCAGGAT" inside the 3' end of AR cDNA was mutated to "TACAAACGTAAATAGTCGAAT" (the protein sequence was maintained the same). For the rescue experiments, doxycycline-inducible AR-overexpressing LNCaP cell lines were infected by either control or AR shRNA lentivirus and selected with 1 µg/ml puromycin and collected for experiments within 5 days (first two days in regular medium and last three days in stripping media before hormone treatment). 20 ng/ml doxycycline was added to the overexpressing samples for 2 days before the 2 hours of vehicle or DHT treatment and collection for experiments including RT-qPCR, Western Blots, and ATAC-seq.

**AR cDNAs for wildtype and mutants**

The full-length human AR cDNA or protein sequences for wildtype and mutants used in this study are below (all synthesized by GenScript) and all are resistant to AR shRNA TRCN0000003718:

ARwt (wildtype): The cDNA sequence for wildtype AR is from NM_000044, the AR protein sequence has 920 amino acids and it matches with UniProt P10275-1.

ARΔLBD mutant: we deleted the 672-920Aa region from the AR wildtype to get the ARΔLBD mutant (with TGA stop code at the end of the cDNA).

ARΔIDR mutant: we deleted the 1-538Aa region from the AR wildtype to get the ARΔIDR mutant. We have added the ATG start code to the cDNA of ARΔIDR for all expression experiments.

AR7FS mutant: For this mutant, we have replaced all 7 phenylalanine (F) amino acids in AR 1-538Aa region with serine (S). The locations of these 7 phenylalanine (F) amino acids are F23, F27, F171, F312, F367, F439, and F497.

ARpQ69 mutant: For this polyQ expansion mutation, we have added additional 46 glutamine (Q) right after the wildtype's original 23 polyQ track (58-80Aa location) to get a pQ69 mutant.

ERαIDR-ARΔIDR mutant: For this swapping mutant, we replaced the AR 1-538Aa IDR region with the IDR region (1-185Aa) from ERα protein (UniProt P03372-1).

FUSIDR-ARΔIDR mutant: For this swapping mutant, we replaced the AR 1-538Aa IDR region with the IDR region (1-214Aa) from FUS protein (UniProt P35637-1).

TAF15IDR-ARΔIDR mutant: For this swapping mutant, we replaced the AR 1-538Aa IDR region with the IDR region (1-205Aa) from TAF15 protein (UniProt Q92804-1).

**Western blotting assay**

Whole cell lysates or nuclear fractions were extracted for Western blotting assays. Cells or nuclei were lysed in RIPA lysis buffer (50 mM Tris-Cl pH 8.0, 150 mM NaCl, NP-40, 0.5% sodium deoxycholate, 0.1% SDS) supplemented with 1 mM DTT, 1 mM PMSF, and 1x protease inhibitor cocktail (Roche). Protein concentrations were quantified with the Bio-Rad protein assay kit. Western blotting was performed as previously described (Liu et al., 2014). Briefly, 30 μg of protein extracts were loaded and separated by SDS–PAGE gels. Blotting was performed with standard protocols using PVDF membrane (Bio-Rad). Membranes were blocked for 1 hour in blocking buffer (5% Non-fat milk in PBST) and probed with primary antibodies at 4°C overnight. After three washes with PBST, the membranes were incubated with HRP-conjugated secondary antibody. Signals were visualized with Clarity Western ECL Substrate (Bio-Rad) as described by the manufacturer. The antibodies used in this assay were: anti-AR (sc-815 for recognizing the C-terminus of AR and sc-7305 for recognizing the N-terminus of AR, Santa Cruz), anti-ERα (sc-8005 for recognizing 2-185Aa region of N-terminal of ERα, Santa Cruz), anti-FUS (sc-373698 for recognizing 2-27Aa region of N-terminal of FUS, Santa Cruz), anti-TAF15 (PA5-84610 for recognizing 55-105Aa region of N-terminal of TAF15, Invitrogen), anti-Histone H3 (A01502, GenScript), anti-Flag (F1804, Sigma), anti-Myc (2276S, Cell Signaling Technology), and anti-GAPDH (sc-25778, Santa Cruz). For the BioID system, the *in vivo* proximity biotinylation events mediated by BioID-tagged AR can be detected by Western blot using streptavidin-HRP (016-030-084, Jackson ImmunoResearch).

**RNA isolation and quantitative RT-PCR**

Total RNA was isolated with RNeasy Mini Kit (Qiagen) according to the manufacturer's protocol and 1μg RNA was used to convert to cDNA using iScript Select cDNA Synthesis Kit (BioRAD) in the presence of both oligo (dT) and random primers. qPCR was conducted with SsoAdvanced Universal SYBR Green Supermix (BioRAD) using CFX384 Real-Time PCR Detection System (BioRAD) according to the manufacturer's instructions. We performed qPCR to test DHT-induced gene activation from gene bodies and also for enhancer activation by testing the levels of eRNAs that are transcribed from enhancer regions. Relative expression of RNAs was determined by the ΔΔCT method using GAPDH as an internal control for quantification analyses of gene targets. Primer sets used in qPCR are as follows: *GAPDH*: F: ACATCATCCCTGCCTCTACTGG, R: GTTTTTCTAGACGGCAGGTCAGG; *AR*: F: TGTCCATCTTGTCGTCTTCG, R: GCTTCTGGGTTGTCTCCTCA; *KLK2*: F: AGCCTGCCAAGATCACAGAT, R: GCAAGAACTCCTCTGGTTCG; *NKX3-1*: F: GCCAAGAACCTCAAGCTCAC, R: AGAAGGCCTCCTCTTTCAGG; *TMPRSS2*: F: CTGGTGGCTGATAGGGGAT, R: GTCTGCCCTCATTTGTCGAT; *NKX3-1* enhancer: F: ACGTAGAAGCCTCTGCTCTAAA, R: GCAATCTTTGGGCCCTGTAATA; *TMPRSS2* enhancer: F: GATGATGAGGTCGTCACAGG, R: CTTCCACGTGCATCTCACAC.

**Luciferase assay**

293T cells were seeded at 20,000 cells per well to 96-well white plate with a transparent bottom (Corning) that had been pre-coated with Poly-D-lysine (Sigma). After overnight, cells were transfected using lipofectamine 2000 (Life Technologies). A DNA mixture was prepared with 600 ng of pEGFP-C3-AR, 200 ng of pCMV-Renilla-3XARE-TATA-luciferase (this construct was generated through gateway LR recombination between entry clone containing 3xARE sequence cloned from Addgene #132360 and destination vector Addgene #101139) and diluted with 100 ul Opti-MEM (Gibco) for transfection of 8 wells of 96-well plate. At the same time, 2 ul of Lipofectamine 2000 was diluted in 100 ul Opti-MEM for transfection of 8 wells of the 96-well plate. After 5 minutes, the diluted Lipofectamine 2000 was mixed with the DNA mixture. During the 20-minute incubation of DNA-lipo transfection mix, cells were washed twice with 100 ul DPBS (Gibco) and incubated with 50 ul of DMEM stripping media. Past the incubation time, 25 ul of the transfection mix was added to each well of the 96-well plate seeded with 293T cells (8 wells for each AR transfection) and incubated for 24 hours. Then cells had the transfection media removed and were treated in quadruplicates with 100 nM EtOH and 100 nM DHT for 24 hours. 48 hours past transfection, firefly and renilla signals were quantified using the Dual-Glo assay kit (Promega) according to its instructions and measured by Cytation 5 machine. The ratio of FLuc/RLuc for each well was calculated and plotted by the average of treatments.

**BioID system setup and pulldown experiment**

The BioID system has been described in our previous publications (Bi et al., 2020; Zhu et al., 2019). The AR cDNAs (WT, ΔIDR, and pQ69) were cloned into pRetroX-mycBioID-MCS at the Not I and Mlu I sites by Gibson reaction (NEB E2611L) to get pRetroX-mycBioID-AR. pRetroX-mycBioID-AR for WT, ΔIDR, and pQ69 were co-transfected with pCL-Ampho packaging plasmid into 293T cell line to produce retroviruses. Retroviruses were used to transduce a parental LNCaP stable line that was engineered to stably express a Tet Repressor using a retroviral vector. G418 (300 μg/ml) and puromycin (0.3 μg/ml) were used for stable selection. Multiple stable cell lines for each BioID transduction were isolated, treated with doxycycline, and screened for BioID-tagged protein expression that was similar to the levels of the respective endogenous genes as revealed by immunoblotting with AR antibody.

The established BioID stable lines were transduced with AR shRNA TRCN0000003718 to knock down the endogenous AR expression level, which will prevent the homodimerization between endogenous wildtype AR protein with overexpressed AR mutants (for a background negative control, we used the LNCaP TetR parental stable line and transduced it with control shRNA to confirm the knockdown efficiency of other lines). All BioID stable cell lines that originally grew in regular medium were washed with PBS three times and started stripping three days before DHT stimulation. To induce mycBioID-AR protein expression, 2 μg/ml doxycycline was added into stripping culture media approximately 24 hours before DHT treatment for 1 hour, followed by incubation with 50mM Biotin for another 24 hours to label all proteins in the proximity of AR before collection.

To identify AR-associated cofactors on chromatin, we purified nuclear fraction for analyses of biotin-labeled proteins. Cytoplasmic and nuclear lysates were isolated as previously described with slight modification (Gagnon et al., 2014). Briefly, cells were scraped from 15 cm dish plates and washed with cold PBS. For every 70 mg of cells, 1 mL of cold Hypotonic Lysis Buffer (HLB) (10 mM Tris-HCl pH 7.5, 3 mM $MgCl_2$, 10 mM NaCl, 0.3% NP-40, 10% glycerol, and 1x protease inhibitor) was used to resuspend the cells followed by the incubation on ice for 10 minutes. Cells were then centrifuged at 800 g for 8 minutes at 4°C. The supernatant was collected as cytoplasmic lysate fraction. The precipitated nuclei were washed 4 times with HLB by pipetting and centrifuging at 200 g for 2 minutes at 4 °C. After HLB washes, the nuclei were resuspended in 0.5 mL of ice-cold Modified RIPA buffer (MRB) (50 mM Tris-HCl pH 7.4, 0.1% SDS, 0.5% sodium deoxycholate, 1% NP-40, 6 mM $MgCl_2$, 150 mM NaCl, 1 mM EDTA, 1 mM DTT, and 1x protease inhibitor). The nuclei pellet in MRB was sonicated till the solution turned clear. 25 U/ml Benzonase and 4 U/ml DNase I were added to the solution and incubated at RT for 30 minutes to digest the chromatin. The solution was then centrifuged at 18,000 g for 15 minutes at 4°C to pellet insoluble protein and other components. The supernatant was collected as the nuclear lysate fraction for the pulldown experiments. For each sample, a small aliquot for each fraction was kept for fractioning result testing using GAPDH and Histone H3 western blots before proceeding with BioID pulldown experiments below.

For BioID pulldown, each nuclear lysate fraction was incubated with 200 µl PBS-washed MyOne Streptavidin T1 Dynabeads (Life Technologies) with overnight rotation at 4°C to pull down all biotinylated proteins in the nuclear lysate. On Day two, the beads were washed 5 times with RIPA buffer (50 mM Tris PH 7.4, 0.4% SDS, 0.1% Triton X-100, 1% NP-40, 300 mM NaCl, 1 mM EDTA, 0.5% sodium deoxycholate, 1 mM DTT) with 10 minutes rotation for each wash (Note: only the first two washes need to use the buffer containing protease inhibitor and be incubated at 4°C. The last three washes can be done at RT without protease inhibitor). After 5X washes with RIPA, the beads were then washed twice more with stringent RIPA buffer containing 5% SDS and then 4X washes with PBS to get rid of SDS residue that might affect trypsin digestion during mass spectrometry. For the $4^{th}$ wash, only 200 µl PBS was used. 10% (20 µl) PBS with beads was transferred to another tube for western blot tests. 90% (180 µl) PBS with beads was kept in the original tube for mass spectrometry. Both tubes were placed on a magnetic stand for 3 minutes, and then all PBS was removed. For western blot tests, the beads were boiled 10 minutes in 2x protein loading buffer with 5% β-Me to release proteins. For mass spectrometry, beads were stored at -80°C before shipping to the Proteomics Core at Sanford-Burnham-Prebys Medical Discovery Institute.

**Mass spectrometry for BioID pulldown beads**

Following immunoprecipitation and washes, proteins were digested directly on the beads. Proteins bound to the beads were resuspended with 8 M urea, 50 mM ammonium bicarbonate, and cysteine disulfide bonds were reduced with 5 mM tris(2-carboxyethyl)phosphine (TCEP) at 30°C for 60 minutes followed by cysteine alkylation with 15 mM iodoacetamide (IAA) in the dark at RT for 30 minutes. Following alkylation, urea was diluted to 1 M using 50 mM ammonium bicarbonate, and proteins were finally subjected to overnight digestion with mass spec grade Trypsin/Lys-C mix (Promega). Finally, beads were pulled down and the solution with peptides was collected into a fresh tube. The beads were then washed once with 50 mM ammonium bicarbonate to increase peptide recovery. Following overnight digestion, samples were acidified with formic acid (FA) and subsequently desalted using AssayMap C18 cartridges mounted on an Agilent AssayMap BRAVO liquid handling system. C18 cartridges were first conditioned with 100% acetonitrile (ACN), followed by 0.1% FA. Samples were then loaded onto the conditioned C18 cartridge, washed with 0.1% FA, and eluted with 60% CAN and 0.1% FA. Finally, the organic solvent was removed in a SpeedVac concentrator prior to LC-MS/MS analysis. Before injecting in the LC-MS, total sample peptide amount was determined by Pierce Quantitative Colorimetric Peptide Assay (ThermoFisher).

Dried samples were reconstituted with 2% acetonitrile and 0.1% formic acid and then analyzed by LC-MS/MS using a Proxeon EASY nanoLC system (ThermoFisher) coupled to Elite mass spectrometer (ThermoFisher). Peptides were separated using an analytical C18 Acclaim PepMap column 75 µm x 500 mm, 2 µm particles (ThermoFisher) in 121 minutes at a flow rate of 300 nL/minute: 1% to 6% B in 1 minute, 6% to 23% B in 56 minutes, 23% to 34% B in 37 minutes, 34% to 48% B in 26 minutes, and 48% to 98% B in 1 minute (A = Formic acid 0.1%; B = 80% ACN: 0.1% Formic acid).

The mass spectrometer was operated in positive data-dependent acquisition mode. MS1 spectra were measured in the Orbitrap with a resolution of 60,000 (AGC target of 1e6, and a mass range from 350 to 1450 m/z). Up to 10 MS2 spectra per duty cycle were triggered, CID-fragmented, and acquired in the linear ion trap (AGC target of 1e4, isolation window of 2 m/z, and a normalized collision energy of 35). Dynamic exclusion was enabled with a duration of 30 seconds.

MS raw files were analyzed by MaxQuant (Cox and Mann, 2008) software with most default settings, and Andromeda (Cox et al., 2011) search engine was used to search against the human Uniprot database. The false discovery rate was set to 0.01 for proteins and peptides with a minimum length of seven amino acids. Variable modifications were set to methionine oxidation and N-terminal acetylation, while fixed modification was set to carbamidomethylation. For label-free protein quantification (Cox et al., 2014), the minimum ratio count was set to two and peptides for quantification was set to unique and razor. Statistical analysis was performed by Perseus(Tyanova et al., 2016). Proteins identified only by site, reverse hits or potential contaminants were removed before downstream analysis. LFQ intensities were log2 transformed, and the matrix was then grouped based on different genetic conditions. The proteins were then filtered by requiring at least two valid values in at least one cell line. Missing values were imputed based on a normal distribution. All the proteins with reduced MS/MS counts in both replicates of ΔIDR or pQ69 AR compared to wildtype AR were uploaded to Enrichr for gene ontology enrichment analyses (Xie et al., 2021).

**ATAC-seq**

ATAC-seq library prep was performed as previously described(Buenrostro et al., 2015). Briefly, 50,000 cells were washed three times with cold PBS, collected by centrifugation then lysed in lysis buffer (10 mM Tris-HCl, pH 7.4, 10 mM NaCl, 3 mM MgCl2, 0.1% NP-40). After purification of nuclei, transposition was performed with Tn5 transposase from Nextera DNA Library Prep Kit (Illumina, catalog # FC-121-1030). Purified DNA was then ligated with adapters, amplified and size selected for sequencing. Libraries were sequenced with Mid 75 bp PE on Illumina NextSeq 500 (4 samples in one lane).

**Global run-on sequencing (GRO-seq)**

To investigate the effect of chemical disruption of AR condensation on androgen-induced transcription, we performed GRO-seq in LNCaP. LNCaP cells were cultured in stripping media for 3 days and then treated with ethanol vehicle control, 100 nM DHT, or 100 nM DHT+2.5% 1,6HD for 2 hours before collection for GRO-seq. GRO-seq experiments were performed as previously reported (Liu et al., 2014). Briefly, ~5-10 million LNCaP cells treated with three different conditions (vehicle, DHT, DHT+1,6HD) were washed 3 times with cold PBS and then sequentially swelled in swelling buffer (10 mM Tris-HCl pH7.5, 2 mM $MgCl_2$, 3 mM $CaCl_2$) for 10 minutes on ice, harvested, and lysed in lysis buffer (swelling buffer plus 0.5% NP-40, 20 units of SUPERase-In, and 10% glycerol). The resultant nuclei were washed two more times with 10 ml lysis buffer and finally re-suspended in 100 µl of freezing buffer (50 mM Tris-HCl pH8.3, 40% glycerol, 5 mM $MgCl_2$, 0.1 mM EDTA). For the run-on assay, re-suspended nuclei were mixed with an equal volume of reaction buffer (10 mM Tris-HCl pH 8.0, 5 mM $MgCl_2$, 1 mM DTT, 300 mM KCl, 20 units of SUPERase-In, 1% sarkosyl, 500 µM ATP, GTP, and Br-UTP, 2 µM CTP) and incubated for 5 minutes at 30°C. The resultant nuclear-run-on RNA (NRO-RNA) was then extracted with TRIzol LS reagent (Life Technologies) following the manufacturer's instructions. NRO-RNA was fragmented to ~300-500 nt by alkaline base hydrolysis on ice for 30 minutes followed by treatment with DNase I and antarctic phosphatase. At this step, only a small portion of all the RNA species was Br-UTP-labeled. To purify the Br-UTP labeled nascent RNA, the fragmented NRO-RNA was immunoprecipitated with anti-BrdU agarose beads (Santa Cruz Biotechnology) in binding buffer (0.5xSSPE, 1 mM EDTA, 0.05% tween) for 1-3 hours at 4°C with rotation. Subsequently, T4 PNK was used to repair the ends of the immunoprecipitated Br-UTP labeled nascent RNA at 37°C for 1 hour. The RNA was extracted and precipitated using acidic phenol-chloroform.

The RNA fragments were subjected to poly-A tailing reaction by poly-A polymerase (NEB) for 30 minutes at 37°C. Subsequently, reverse transcription was performed using oNTI223 primer and SuperScript III RT kit (Life Technologies). The cDNA products were separated on a 10% polyacrylamide TBE-urea gel, and only those fragments migrating between 100-500 bp were excised and recovered by gel extraction. Next, the first-strand cDNA was circularized by CircLigase (Epicentre) and re-linearized by APE1 (NEB). Re-linearized single-stranded cDNA (sscDNA) was separated on a 10% polyacrylamide TBE gel and the appropriately sized product (~120-320 bp) was excised and gel-extracted. Finally, the sscDNA template was amplified by PCR using the Phusion High-Fidelity enzyme (NEB) according to the manufacturer's instructions. The oligonucleotide primers oNTI200 and oNTI201 were used to generate DNA libraries for deep sequencing. The sequences for primers oNTI223, oNTI200, and oNTI201 are as follows:
oNTI223-TruSeqLT:
/5Phos/AGATCGGAAGAGCGTCGTGTA/idSp/CAGACGTGTGCTCTTCCGATC TTTT TTT TTT TTT TTT TTT TVN
oNTI201-TruseqLT:
5'AATGATACGGCGACCACCGAGATCTACACTCTTTCCCTACACGACGCTCTTCCGATCT 3'
oNTI200-TruSeqID1:
5'CAAGCAGAAGACGGCATACGAGAT CGTGAT GTGACTGGAGTT CAGACGTGTGCTCTTCCGATC 3'

**Establishing stable cell lines expressing BLRP-Tagged AR and biotin ChIP-seq**

To study genome-wide binding patterns for wildtype and mutants of AR, we have established a parental LNCaP stable cell line that expressed BirA enzyme and Tet-Repressor similar to what we have done previously in MCF7 cells (Liu et al., 2014). We then used this parental cell line to establish doxycycline-inducible stable cell lines expressing BLRP-tagged wildtype or mutant AR proteins (the cDNAs are resistant to AR shRNA TRCN0000003718) at close to endogenous levels. For each line, we have prepared two different cell samples for biotin ChIP: the control one was transduced with control shRNA and not treated with doxycycline; the experimental sample was transduced with AR shRNA TRCN0000003718 and treated by 2 µg/ml doxycycline for 24 hours. Both cell samples were hormone-stripped in phenol red-free RPMI1640 medium plus 5% charcoal-treated FBS for 3 days, followed by treatment with 100 nM DHT for 2 hours before collection for biotin ChIP experiments. Biotin ChIP experiments for BLRP-tagged wildtype or mutant AR were performed with our previously published protocol (Liu et al., 2014). Briefly, cross-linked protein-DNA complexes were pulled down by Nanolink Streptavidin Magnetic beads (Solulink) and the washing was performed under much more stringent conditions that included 4 washes with 1% SDS in TE (20 minutes each) and two washes with 1% Triton X-100 in TE. The washed streptavidin beads were then subjected to AcTEV protease (Life Technologies) digestion twice for tagged protein-DNA complex elution before de-crosslinking at 65°C overnight. After de-crosslinking overnight at 65°C, the final ChIP DNA was extracted and purified using QIAquick spin columns (Qiagen). BLRP-tagged AR inducible cell lines without doxycycline induction were used as ChIP-seq control for background detection.

The ChIP-seq libraries were constructed using KAPA's HyperPrep kit (KK8504), followed by deep sequencing with Illumina's HiSeq 3000 system according to the manufacturer's instructions.

**Statistical analysis**

Each experiment was repeated at least twice to make sure that the results are reproducible. Results are representative of at least three biological replicates and reported as mean ± SEM or mean ± SD. Data were analyzed and statistics were performed using unpaired two-tailed Student's t-tests or one-way ANOVA (Prism 5 GraphPad). Significant differences between two groups were noted by asterisks (* $P<0.05$, ** $P<0.01$, *** $P<0.001$).

**Structure prediction using AlphaFold**

The AlphaFold2 algorithm was employed locally (Jumper et al., 2021). The full-length AR protein sequences were used to predict their structures. Casp14 preset was selected and the predicted structure with the highest confidence after Amber relaxation was used for protein structure visualization.

**ChIP-seq data analysis**

We retrieved publicly available AR and H3K27ac ChIP-seq data from GSE83860 and GSE51621 respectively. Reads were aligned to the human genome (hg19) using bowtie with "--best --strata –m 1" parameters (Langmead et al.). Duplicated reads were eliminated for subsequent analysis. MACS2 was employed to call peaks comparing immunoprecipitated chromatin with input chromatin using standard parameters and a q-value cutoff of 1e-5 (Zhang et al.). The peaks overlapping with the blacklist regions from UCSC were removed. The H3K27ac peaks located 10kb away from transcriptional start sites of annotated genes were defined as enhancers, and those co-localized with AR peaks were defined as AR-bound enhancers. Since the most active AR-bound enhancer regions usually generate strong eRNA signals upon DHT treatment, we further derived 1,158 active AR enhancers based on the induction of eRNA from GRO-seq data comparing DHT with vehicle conditions. 1,500 randomly selected non-AR bound enhancers were used as control.

For Biotin ChIP-seq results, reads were aligned to the human genome (hg19) using bowtie v1.1.2 with "--best --strata –m 1" parameters (Langmead et al., 2009). Only uniquely mapped and without-duplication reads were selected for subsequent analysis. MACS2 was employed to call peaks by comparing immunoprecipitated chromatin with input chromatin using standard parameters and a q-value cutoff of 1e-5 (Zhang et al., 2008). The peaks overlapping with the blacklist regions downloaded from UCSC were removed. ChIPseeker was used to annotate the peaks (Yu et al., 2015).

For all the aggregation plots, the signal fold change (FC) was calculated as the ratio of average tag density. The library-size normalized tag density within a 1 kb region around the peak center (+/- 500 bp) from each condition was used for calculation. P-value was determined by Wilcoxon signed-rank test.

**ATAC-seq data analysis**

ATAC-seq reads were adapter trimmed using cutadapt 1.11, and aligned to the human genome (hg19) using bowtie with parameters "--best --strata –m 1 –v 2" (Martin). Aligned reads with the same genomic position and orientation were collapsed to a single read. The reads were extended to 200bp and normalized to a sequencing depth of ten million reads for each library.

**GRO-seq data analysis**

GRO-seq reads were aligned to the human genome (hg19) using bowtie with "--best --strata –m 1 –v 2" parameters. Duplicated reads were eliminated for subsequent analysis. To balance the clonal amplification bias and total useful reads, only three reads at most were allowed for each unique genomic position. When measuring the expression level of genes, mapped reads from the first 30 kb of the gene body were counted, excluding promoter-proximal region (transcription start site (TSS) to 1000 bp downstream of TSS; if the length of a gene is shorter than 10kb, then the reads mapped to first 10%*length regions were excluded). If the length of a gene is shorter than 30kb, then mapped reads from the whole gene were counted, excluding promoter-proximal region and gene end (500 bp upstream of transcription termination site (TTS) to TTS).

**Data visualization**

All ChIP-seq, ATAC-seq, and GRO-seq data was visualized in Integrative Genomics Viewer (IGV) (Robinson et al.). For GRO-seq, the reads were separated by strand and extended to a length of 100 nt in the 5'-to'3' direction; for ChIP-seq and ATAC-seq, the reads were extended to 200 nt in the 5'-to-3' direction. All data were normalized to 10 million uniquely mapped reads per experiment. A customized R script was used to generate heatmap and line plots for ChIP-seq, ATAC-seq, and GRO-seq data.

**DATA AND SOFTWARE AVAILABILITY**

The GEO accession number for all deep sequencing data reported in this paper is GSE215163, which includes the ChIP-seq data set (GSE215161), ATAC-seq data set (GSE215160), and GRO-seq data set (GSE215162).

We also used some published ChIP-seq data from the Gene Expression Omnibus database for AR and H3K27ac under accession number GSE83860 (Malinen et al., 2017) and GSE51621 (Hazelett et al., 2014) respectively.

### METHODS REFERENCE

Bi, M., Zhang, Z., Jiang, Y.Z., Xue, P., Wang, H., Lai, Z., Fu, X., De Angelis, C., Gong, Y., Gao, Z.*, et al.* (2020). Enhancer reprogramming driven by high-order assemblies of transcription factors promotes phenotypic plasticity and breast cancer endocrine resistance. Nat Cell Biol *22*, 701-715.

Buenrostro, J.D., Wu, B., Chang, H.Y., and Greenleaf, W.J. (2015). ATAC-seq: A Method for Assaying Chromatin Accessibility Genome-Wide. Curr Protoc Mol Biol *109*, 21 29 21-29.

Chong, S., Dugast-Darzacq, C., Liu, Z., Dong, P., Dailey, G.M., Cattoglio, C., Heckert, A., Banala, S., Lavis, L., Darzacq, X.*, et al.* (2018). Imaging dynamic and selective low-complexity domain interactions that control gene transcription. Science *361*.

Cox, J., Hein, M.Y., Luber, C.A., Paron, I., Nagaraj, N., and Mann, M. (2014). Accurate proteome-wide label-free quantification by delayed normalization and maximal peptide ratio extraction, termed MaxLFQ. Mol Cell Proteomics *13*, 2513-2526.

Cox, J., and Mann, M. (2008). MaxQuant enables high peptide identification rates, individualized p.p.b.-range mass accuracies and proteome-wide protein quantification. Nat Biotechnol *26*, 1367-1372.

Cox, J., Neuhauser, N., Michalski, A., Scheltema, R.A., Olsen, J.V., and Mann, M. (2011). Andromeda: a peptide search engine integrated into the MaxQuant environment. J Proteome Res *10*, 1794-1805.

Gagnon, K.T., Li, L., Janowski, B.A., and Corey, D.R. (2014). Analysis of nuclear RNA interference in human cells by subcellular fractionation and Argonaute loading. Nat Protoc *9*, 2045-2060.

Hazelett, D.J., Rhie, S.K., Gaddis, M., Yan, C., Lakeland, D.L., Coetzee, S.G., Ellipse, G.-O.N.c., Practical, c., Henderson, B.E., Noushmehr, H.*, et al.* (2014). Comprehensive functional annotation of 77 prostate cancer risk loci. PLoS Genet *10*, e1004102.

Janicki, S.M., Tsukamoto, T., Salghetti, S.E., Tansey, W.P., Sachidanandam, R., Prasanth, K.V., Ried, T., Shav-Tal, Y., Bertrand, E., Singer, R.H.*, et al.* (2004). From silencing to gene expression: real-time analysis in single cells. Cell *116*, 683-698.

Jumper, J., Evans, R., Pritzel, A., Green, T., Figurnov, M., Ronneberger, O., Tunyasuvunakool, K., Bates, R., Zidek, A., Potapenko, A.*, et al.* (2021). Highly accurate protein structure prediction with AlphaFold. Nature *596*, 583-589.

Langmead, B., Trapnell, C., Pop, M., and Salzberg, S.L. (2009). Ultrafast and memory-efficient alignment of short DNA sequences to the human genome. Genome Biol *10*, R25.

Liu, Z., Merkurjev, D., Yang, F., Li, W., Oh, S., Friedman, M.J., Song, X., Zhang, F., Ma, Q., Ohgi, K.A.*, et al.* (2014). Enhancer activation requires trans-recruitment of a mega transcription factor complex. Cell *159*, 358-373.

Malinen, M., Niskanen, E.A., Kaikkonen, M.U., and Palvimo, J.J. (2017). Crosstalk between androgen and pro-inflammatory signaling remodels androgen receptor and NF-kappaB cistrome to reprogram the prostate cancer cell transcriptome. Nucleic Acids Res *45*, 619-630.

Martin, M. (2010). Cutadapt removes adapter sequences from high-throughput sequencing reads. EMBnet *17*, 10-12.

Robinson, J.T., Thorvaldsdottir, H., Winckler, W., Guttman, M., Lander, E.S., Getz, G., and Mesirov, J.P. (2011). Integrative genomics viewer. Nat Biotechnol *29*, 24-26.

Sabari, B.R., Dall'Agnese, A., Boija, A., Klein, I.A., Coffey, E.L., Shrinivas, K., Abraham, B.J., Hannett, N.M., Zamudio, A.V., Manteiga, J.C.*, et al.* (2018). Coactivator condensation at super-enhancers links phase separation and gene control. Science *361*.

Shin, Y., Berry, J., Pannucci, N., Haataja, M.P., Toettcher, J.E., and Brangwynne, C.P. (2017). Spatiotemporal Control of Intracellular Phase Transitions Using Light-Activated optoDroplets. Cell *168*, 159-171 e114.

Tyanova, S., Temu, T., Sinitcyn, P., Carlson, A., Hein, M.Y., Geiger, T., Mann, M., and Cox, J. (2016). The Perseus computational platform for comprehensive analysis of (prote)omics data. Nat Methods *13*, 731-740.

Xie, Z., Bailey, A., Kuleshov, M.V., Clarke, D.J.B., Evangelista, J.E., Jenkins, S.L., Lachmann, A., Wojciechowicz, M.L., Kropiwnicki, E., Jagodnik, K.M.*, et al.* (2021). Gene Set Knowledge Discovery with Enrichr. Curr Protoc *1*, e90.

Yu, G., Wang, L.G., and He, Q.Y. (2015). ChIPseeker: an R/Bioconductor package for ChIP peak annotation, comparison and visualization. Bioinformatics *31*, 2382-2383.

Zhang, Y., Liu, T., Meyer, C.A., Eeckhoute, J., Johnson, D.S., Bernstein, B.E., Nusbaum, C., Myers, R.M., Brown, M., Li, W.*, et al.* (2008). Model-based analysis of ChIP-Seq (MACS). Genome Biol *9*, R137.

Zhu, C., Li, L., Zhang, Z., Bi, M., Wang, H., Su, W., Hernandez, K., Liu, P., Chen, J., Chen, M.*, et al.* (2019). A Non-canonical Role of YAP/TEAD Is Required for Activation of Estrogen-Regulated Enhancers in Breast Cancer. Mol Cell *75*, 791-806 e798.
